# The shared *neo*antigen landscape of MSI cancers reflects immunoediting during tumor evolution

**DOI:** 10.1101/691469

**Authors:** Alexej Ballhausen, Moritz Jakob Przybilla, Michael Jendrusch, Saskia Haupt, Elisabeth Pfaffendorf, Markus Draxlbauer, Florian Seidler, Sonja Krausert, Aysel Ahadova, Martin Simon Kalteis, Daniel Heid, Johannes Gebert, Maria Bonsack, Sarah Schott, Hendrik Bläker, Toni Seppälä, Jukka-Pekka Mecklin, Sanne Ten Broeke, Maartje Nielsen, Vincent Heuveline, Julia Krzykalla, Axel Benner, Angelika Beate Riemer, Magnus von Knebel Doeberitz, Matthias Kloor

## Abstract

The immune system can recognize and attack cancer cells, especially those with a high load of mutation-induced *neo*antigens. Such *neo*antigens are particularly abundant in DNA mismatch repair (MMR)-deficient, microsatellite-unstable (MSI) cancers. MMR deficiency leads to insertion/deletion (indel) mutations at coding microsatellites (cMS) and to *neo*antigen-inducing translational frameshifts. The abundance of mutational *neo*antigens renders MSI cancers sensitive to immune checkpoint blockade. However, the neoantigen landscape of MMR-deficient cancers has not yet been systematically mapped. In the present study, we used a novel tool to monitor *neo*antigen-inducing indel mutations in MSI colorectal and endometrial cancer. Our results show that MSI cancers share several highly immunogenic *neo*antigens that result from specific, recurrent indel mutation events. Notably, the frequency of such indel mutations was negatively correlated to the predicted immunogenicity of the resulting *neo*antigens. These observations suggest continuous immunoediting of emerging MMR-deficient cells during tumor evolution.

**One sentence summary:** Quantitative indel mutation analysis reveals evidence of immune selection in mismatch repair-deficient cancers

## Main text

DNA MMR deficiency is a major mechanism causing genomic instability in human cancer. MMR-deficient cancers accumulate an exceptionally high number of somatic mutations. These mutations encompass certain types of single nucleotide alterations, but mostly insertion/deletion (indel) mutations at repetitive sequence stretches termed microsatellites (microsatellite instability, MSI) (*1, 2*).

About 15% of colorectal cancers (CRC), up to 30% of endometrial cancers (EC) and multiple other tumors display the MSI phenotype (*3*). MSI tumors can develop sporadically or in the context of Lynch syndrome, the most common inherited cancer predisposition syndrome. Due to this very specific process of genomic instability, the pathogenesis of MSI cancers can be precisely dissected (*4*): Indel mutations affecting coding microsatellites (cMS), predominantly coding mononucleotide repeats, in genomic regions encoding tumor suppressor genes are considered major drivers of MSI tumorigenesis (*5–7*). Importantly, the same indels that inactivate tumor suppressor genes simultaneously cause translational frameshifts, thereby generating unique frameshift peptide (FSP) *neo*antigens (*4, 8*).

For the recognition of *neo*antigens by the immune system, processing through the cellular antigen processing machinery and presentation by human leukocyte antigen (HLA) class I molecules on the tumor cell surface are essential prerequisites. These HLA class I molecules consist of a heavy chain and a non-covalently bound light chain (encoded by the *Beta-2-microglobulin [B2M]* gene), both of which are essential for functional antigen presentation. The likelihood of HLA binding for a defined peptide depends on the HLA genotype, as every individual harbors six alleles (*HLA-A*, *HLA-B*, *HLA-C*, two alleles each) which encode for HLA class I heavy chains (*9*).

The specific mutational steps required for malignant transformation during the evolution of MSI tumors are thus also responsible for their pronounced immunogenicity. MSI tumors are commonly associated with dense lymphocyte infiltration and pronounced local responses of the adaptive immune system at the tumor site (*10–14*). Immune recognition of MSI tumor cells is not only responsible for a comparatively favorable clinical course, but also reflected by the high sensitivity of advanced stage MSI cancers towards immune checkpoint blockade (ICB) (*15, 16*). However, some patients do not respond to ICB treatment.

We and others previously showed that specific T cell responses against a few MMR deficiency-induced frameshift *neo*antigens occur prior to and after immune checkpoint blockade (*15, 17, 18*). However, the landscape of *neo*antigens and potential epitopes in MSI cancer has not been described systematically. One important reason for this gap of knowledge is the fact that short-read next-generation sequencing (NGS) approaches have a limited sensitivity for the detection of indel mutations at homopolymer sequences such as *neo*antigen-related cMS (*19–21*).

Here, we map the FSP *neo*antigen and epitope landscape of the two most common MMR-deficient cancer types, colorectal and endometrial cancer, using a newly developed tool for the quantification of cMS mutation patterns (ReFrame) combined with NetMHCpan 4.0, a state-of-the-art *in silico* epitope prediction tool (*9*). Using ReFrame, we were able to reveal a set of previously unknown shared FSP *neo*antigens in MSI cancers.

Our results provide strong evidence for continuous immunoediting during MSI tumor evolution and underline the potential of *neo*antigen-based vaccines against MSI cancers.

### cMS mutation frequencies in MSI CRC and EC

Short-read NGS approaches are not ideally suited for mutational and *neo*antigen profiling of MSI cancers (*19–21*), showing a high variability in mutation frequency regarding the detection of mutations in different cMS candidates like i.e. TGFBR2, SLC35F5 or TFAM (Table S1). Importantly cMS repeats of increased length which are most susceptible to mutations and therefore encompass the most important mutational targets during MMR-deficient tumorigenesis, are missed by NGS technology that is in common use today (*3, 5, 19, 22–26*). To fill this gap and precisely quantify cMS mutation patterns and their resulting *neo*antigen frames in MMR-deficient cancers, we developed a novel algorithm based on fragment length analysis as the current gold standard for the detection of MSI. ReFrame, our REgression-based FRAMEshift quantification algorithm, allows unbiased quantitative detection of indel mutations by solving a linear system of mathematical equations to remove stutter band artifacts, which result from polymerase slippage events during PCR amplification and subsequent nucleotide gains and losses similar to MSI-induced indels (Fig. S1).

We used ReFrame in a series of MSI colorectal cancers (MSI CRCs; n=139) (Table S2) to screen for mutations in 41 cMS residing in 40 target genes derived from the first comprehensive cMS database (<u>Seltarbase</u>) (*27*). Additionally, we investigated mutation profiles in a cohort of MSI endometrial cancers (MSI ECs; n=14).

In agreement with previous reports (*23, 27*), our results show that the load of indels at cMS in MSI CRC and EC is high and that multiple concomitant indels at several cMS in the same tumor are very common. Although most CRC and EC were distinguishable based on the cMS mutation patterns, a large set of cMS mutations were shared by the majority of MSI CRC and/or MSI EC (Fig. 1, Fig. S2).

**Fig. 1.**
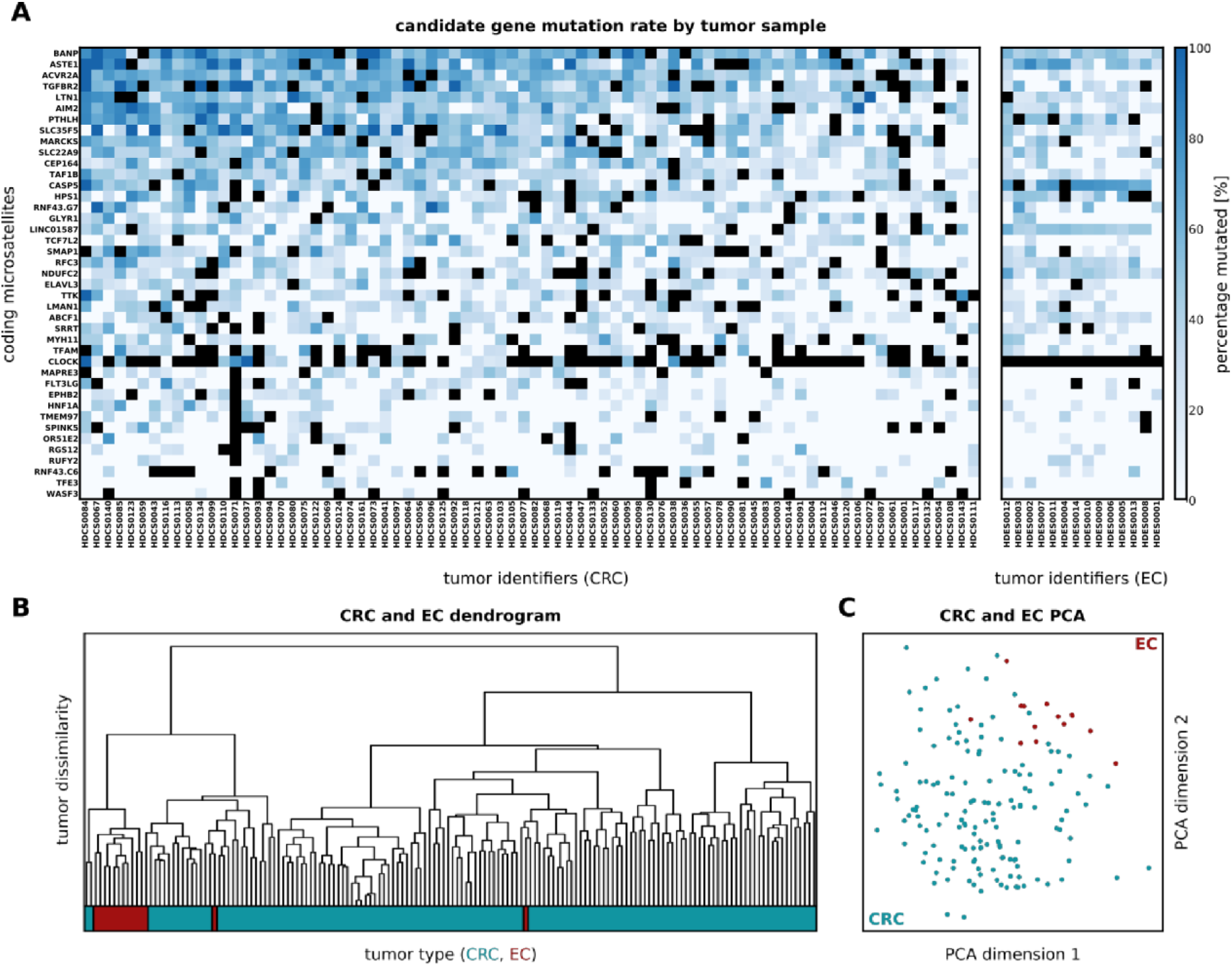
Mutation frequencies of coding microsatellites (cMS) in MSI colorectal (CRC) and endometrium cancer (EC). **(A)** The relative frequency of mutant alleles is shown for 41 cMS (rows) in CRC and EC tumor samples (columns) derived using ReFrame. cMS were sorted top-down according to their mutation frequency indicated by blue boxes of different intensity. Dark blue represents high mutation frequency, whereas pale blue represents low mutation frequency. Black boxes indicate missing data points. The respective tumor samples are shown below each column. cMS were analyzed for both CRC and EC and are depicted separately for each tumor type (left panel: CRC, right panel: EC). The complete dataset can be found in the Supplementary Material. **(B)** Dendrogram of CRC and EC samples with respect to their mutation patterns, where the color bar below indicates the tumor type. **(C)** Principal component analysis of EC (red) and CRC (blue) samples with respect to their mutation patterns. EC samples are distinguishable from CRC samples by mutation patterns alone.

Moreover, we observed a significant variation of the number of mutations per tumor, ranging from 8 to 29 (median=20) out of 41 analyzed cMS in MSI CRC and from 8 to 25 in MSI EC (median=18). The observed variation suggests potential differences in the *neo*antigen load of MSI tumors. Potential clinical consequences, e.g. for the sensitivity towards ICB, should be assessed in future clinical studies (*28–30*).

ReFrame is not only able to quantify mutation frequency, but also to distinguish mutation types, which is crucial for the prediction of the frame of the resulting *neo*antigen. As the translation of nucleotide into amino acid sequences is based on three base codons, every mutation in a homopolymer region can either result in a simple deletion or insertion of amino acids or in two entirely different *neo*antigen reading frames: Deletions of one nucleotide (further referred to as *minus 1* or *m1*) or insertions of two nucleotides (*plus 2*, *p2*) will result in a shift to a frame here referred to as “minus-one” (M1), while deletions of two nucleotides (*minus 2, m2*) or insertion of one nucleotide (*plus 1, p1*) will result in a shift to a frame referred to as “minus-two” (M2) (Fig. 2, Fig. S3, Table S3). The results demonstrate that *m1* mutations, resulting in M1 reading frames, were the predominant mutation type (77% in MSI CRC, Fig. 2). The M1/M2 distribution varied significantly across distinct cMS, with significantly elevated numbers of M2 mutation in BANP, TAF1B and ELAVL3, whereas in ACVR2A, HPS1, SLC35F5 and TCF7L2 there were significantly more mutations leading to an M1 frameshift than expected by chance (Bonferroni corrected binomial test, p<0.05; Table S4 and S5).

**Fig. 2.**
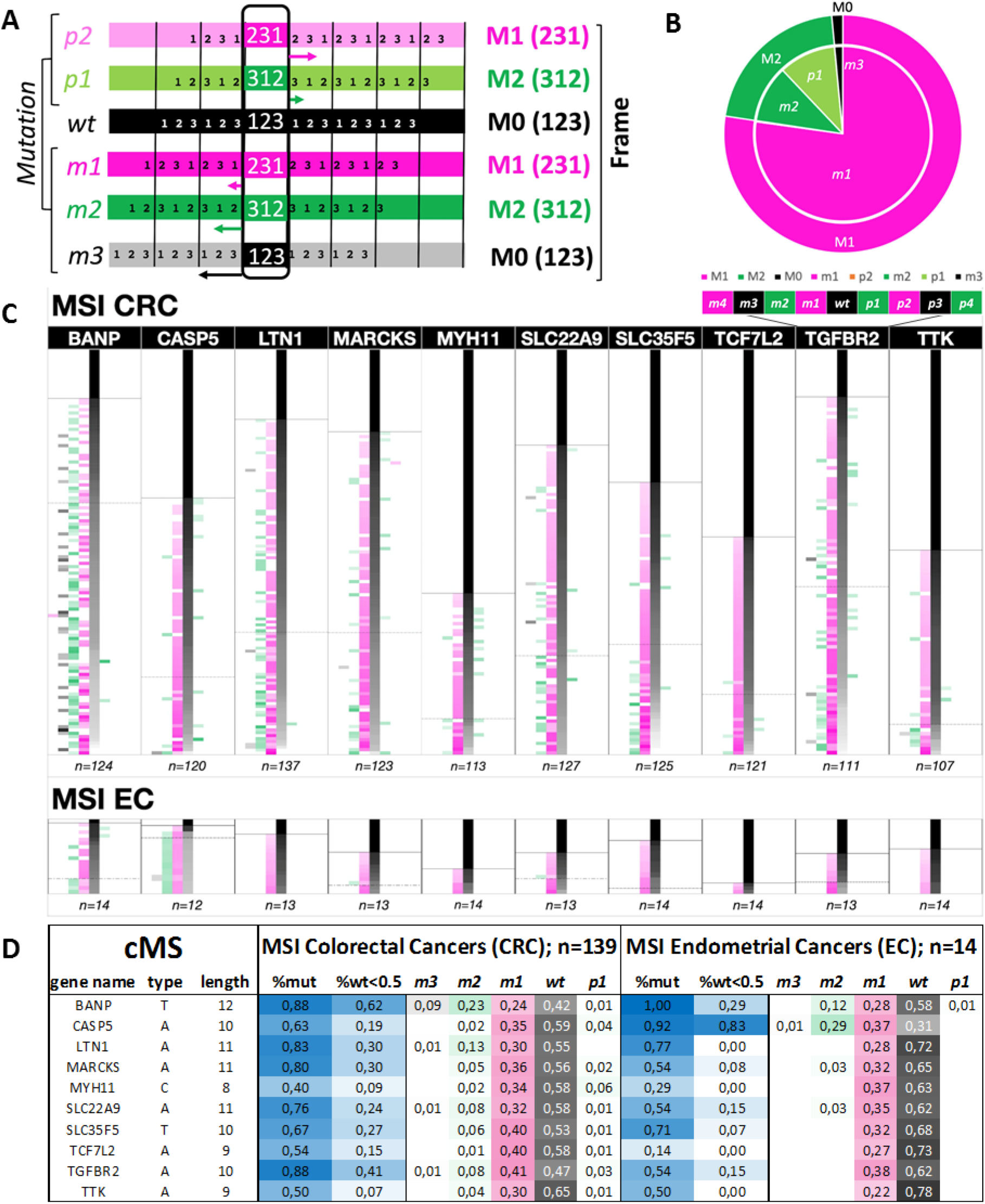
Mutational pattern distribution in cMS based on ReFrame analysis. (A) Scheme of frameshift mutations (*m3* – *p2*) on the left and their corresponding frames (M0, M1, M2) on the right. The numbers 1-3 indicate base triplets in the corresponding frame. Arrows mark the shift between the wt frame and the alternate frame. (B) Distribution of all cMS frameshift mutations (*m3* – *p2*) and their corresponding frames (M0, M1, M2) in MSI CRC quantified using ReFrame. Mutations in all MSI CRC samples were classified in corresponding reading frames (M0, black; M1, magenta; M2, green) and their overall allele ratios quantified. (C) The detailed mutational patterns of 10 representative cMS are depicted with their respective frequency of mutation for all possible resulting frameshift mutations (m4 – p4) in MSI CRC and EC (see Fig. S3 for complete dataset). Each row constitutes one analyzed tumor sample with its related allele ratios. For each cMS, tumors were sorted by the proportion of wild type alleles top to bottom. The number of samples analyzed for a certain candidate is indicated below each candidate’s figure. Color indicates the resulting reading frames: magenta indicates the M1 frame, corresponding to *m1*, *m4* and *p2* mutations; green indicates the M2 frame, corresponding to *m2*, *p1*, *p4* mutations. Because *wt*, *m3* and *p3* mutations do not result in translational frameshifts, they are shown in black (M0) (see also magnification, right panel). Intensities represent ReFrame-calculated ratios from white (0%) to full intensity magenta/green/black (100%) according to the resulting reading frame of the column. The annotated solid lines (first horizontal line top down) show the end of the non-mutated tumor samples while the dotted lines (second horizontal line top down) mark the beginning of tumors being more than 50% mutated, associated with biallelic hits within the respective sample. (D) Calculated mutation frequencies and mean allele ratios of most common mutation types (*m3* – *p1*) resulting from ReFrame analysis in 10 representative cMS (see Tab. S2 for complete dataset). The table is showing an overview of cMS with mutation frequencies above 50% in MSI CRC or MSI EC, depicting the mutation frequencies (%mut), the ratio of samples with biallelic hits, indicated by a proportion of wt alleles lower than 50% of all the detected signals (%wt<0.5), as well as the mean mutational pattern for the cMS candidates sorted by their length. The allele ratios are depicted for wild-type (wt), minus one up to three base pair deletions (*m1* – *m3*) and one base pair insertions as *m4* or *p2* – *p4* mutations only rarely occurred or were completely absent. The same color code as depicted in (C) was used.

### Epitope landscape of MSI *neo*antigens

Following the detection of shared indel mutations in MSI colorectal and endometrial cancers, we evaluated the possible immunogenic potential of the frameshift *neo*antigens and associated *neo*peptides resulting from antigen processing.

We used NetMHCpan 4.0, a state-of-the-art HLA binding prediction tool based on artificial neural networks, to predict *neo*peptides that are possibly presented as epitopes by HLA class I antigens encoded by the most important HLA supertypes (*9, 31, 32*). Applying commonly accepted IC_50_ thresholds we distinguished between three classes of peptides with high (IC_50_ < 50 nM), low (50 nM < IC_50_ < 500 nM) and very low (500 nM < IC_50_ < 5000 nM) predicted HLA binding affinity (*31, 33*). As a first step, we analyzed all possible frameshift *neo*antigen sequences derived from the M1 and M2 frameshifts of the 41 cMS. We then complemented this set to cover all possible FSP *neo*antigens (n=524) derived from 264 cMS with a length of 8 or more nucleotides published in Seltarbase (Data S1) (*27*). Our results indicate multiple FSPs resulting from M1 or M2 frameshift mutations, that are potentially recognized by the immune system. We detected a wide range of variability with regard to the number of predicted putative epitopes maximally contained within a defined *neo*antigen. The highest number of predicted putative high-affinity epitopes within a *neo*antigen was 23 (for the M1 frame of P4HB), (low affinity: 92 predicted putative epitopes in M1 SPINK5; very low-affinity: 375 predicted putative epitopes in M1 P4HB). Other cMS mutation-induced *neo*antigens showed a complete lack of predicted epitopes (Fig. 3, Data S2 and S5).

**Fig. 3.**
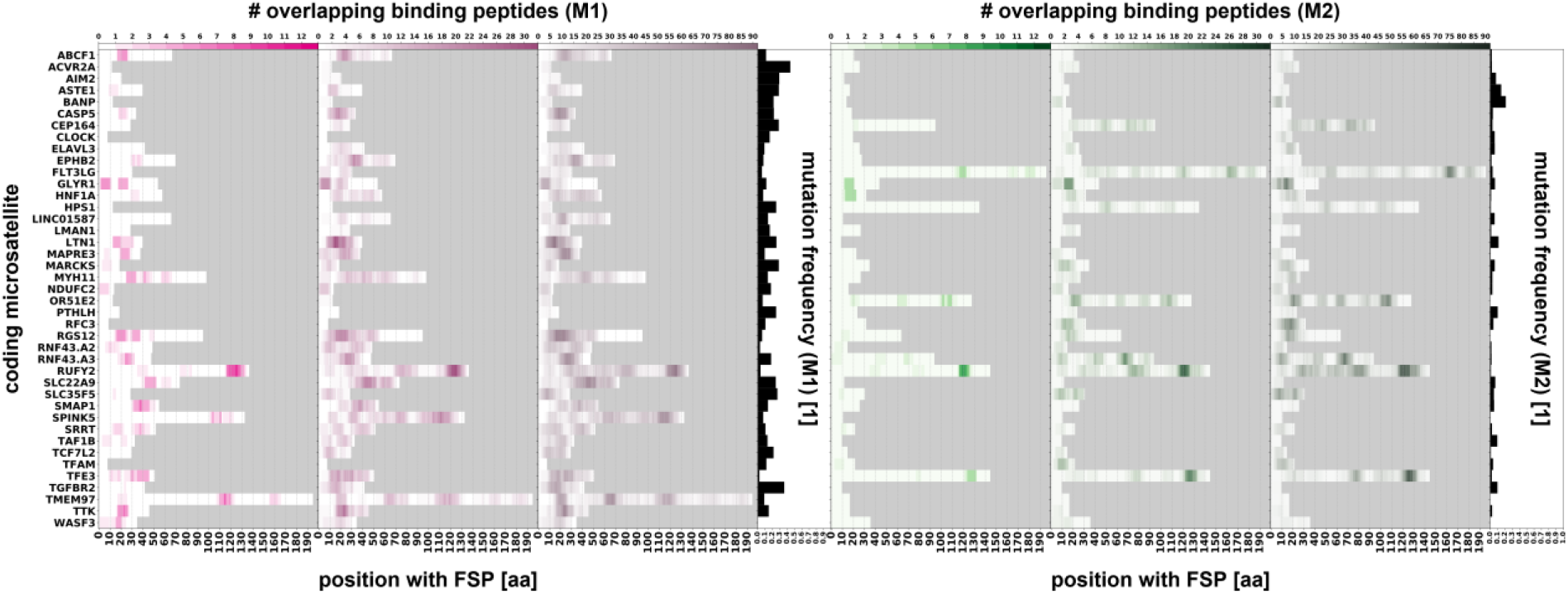
HLA binding predictions for a dataset of 82 analyzed frameshift *neo*antigen peptides. The figures display the predicted epitopes for the M1 and M2 FSP *neo*antigens derived from all 41 cMS candidates over the length of the respective peptide. All epitopes predicted for three binding affinity thresholds (IC_50_ < 50 nM, IC_50_ < 500 nM and IC_50_ < 5000 nM) are shown for the M1 FSPs (left, magenta) and the M2 FSPs (right, green). All candidates are sorted in alphabetical order with their respective frequency of mutation according to the ReFrame results. The grey field is highlighting the ends of the respective FSPs.

For HLA-A*02:01, the most common HLA allele in the USA European Caucasian population (*34*), one or more high-affinity peptides were predicted for 19.8% of the FSP *neo*antigens. HLA-A*02:01 epitopes with lower affinity were present in 39.5% (≤ 500 nM) and 59.8% (≤ 5000 nM) of candidates (Fig. S4, Table S6, Data S3).

To make the potential impact of certain cMS candidates more tangible and to identify frameshift *neo*antigens with potentially highest relevance for immune recognition, we defined a “general epitope likelihood score” (GELS; see method section “Computation of immunological scores”). GELS accounts for HLA binding prediction and the prevalence of the respective HLA allele in a defined population, as the latter influences the probability of a *neo*peptide to be an epitope recognized by the immune system in a patient of this population (*9, 34*). We calculated GELS for all FSP neoantigens using HLA allele frequencies for USA European Caucasians (calculations for additional ethnic groups are provided in Data S3).

Accounting for a potential relation between immunogenicity and mutation frequency, we noticed that the most commonly mutated cMS located in the ACVR2A gene showed a very low GELS (*p*_*mut*_ = 91%, GELS = 5.1%), whereas very high GELS candidates seemed to be associated with a low mutation frequency (i.e. TMEM97, *p*_*mut*_ = 27%, GELS = 91.1%; SPINK5, *p*_*mut*_ = 26%, GELS = 91.1%; RUFY2, *p*_*mut*_ = 16%, GELS = 90.4%; *p*_*binding*_ = 50% in USA European Caucasian population; Data S3). Hierarchical clustering of cMS candidates on all tumor samples revealed the existence of three distinct populations of cMS (Fig. 4A), which was retained in *B2M*-wild type, but not *B2M*-mutant tumors (Fig. 4B).

**Fig. 4.**
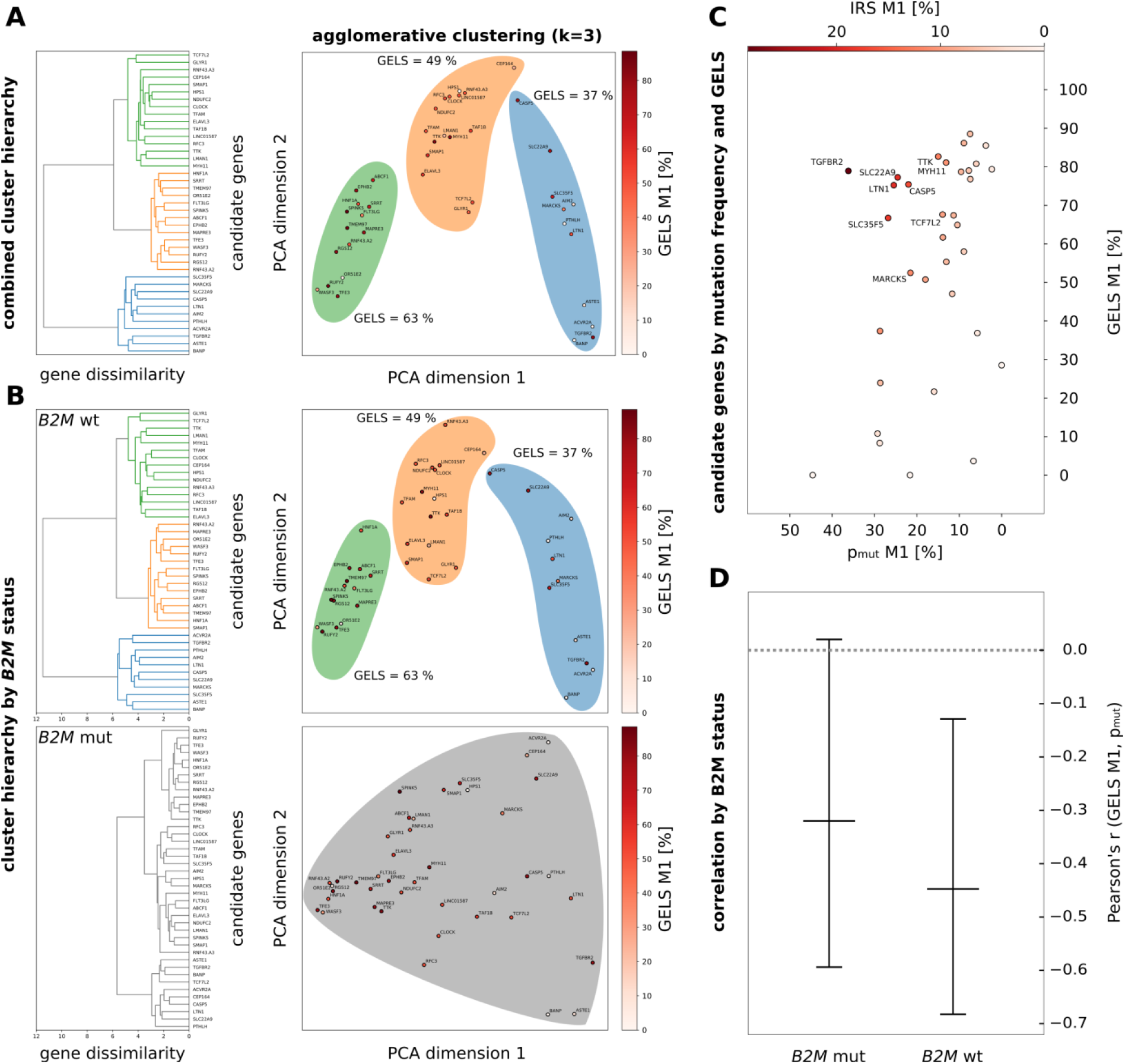
Evidence of immune selection from cMS mutation patterns and general epitope likelihood score (GELS). **(A)** Hierarchical clustering of cMS candidates on all tumor samples using frameshift abundance features. The full clustering hierarchy is displayed as a dendrogram, showing three clusters (blue, yellow, green) of genes. The same clusters are visualized using RBF kernel PCA with two principal components, and colored by their GELS. The three clusters display a trend in their mean GELS, with increasing values from green to blue. **(B)** Hierarchical clustering of cMS candidates on features split by tumor *B2M* status, with wildtype features at the top and mutated features at the bottom. The full hierarchy is displayed as a dendrogram for both feature sets, with the threshold dissimilarity for clustering indicated by a red line. Here, *B2M*-mutated features show no clustering at the given dissimilarity threshold. The same data is again shown using RBF kernel PCA with two principal components. While the wildtype data shows the same clusters as all tumors combined, the clustering is lost in the case of mutated *B2M*. **(C)** The mutational frequency of FSPs resulting from one base pair deletions (m1) is shown on the y axis against the GELS of the resulting M1 FSP neoantigens (x axis). For the calculation of the GELS, all predicted epitopes (IC50 < 500 nM) were taken into account, with an assumed probability for a binder to be a true positive of p_binding_=50%. Every bubble depicted represents one candidate. The gradient intensity of the bubbles shows the IRS, with white color representing a low IRS, while dark red displays a high IRS. All candidates with an IRS of 10% or higher are annotated. **(D)** Correlation between the number of predicted epitopes in cMS mutation-induced FSP neoantigens and the frequency of the respective cMS mutations in MSI colorectal cancer separated by *B2M* mutation status. The Pearson’s r from the correlation test is shown on the y-axis, while the different groups of tumors are shown on the x-axis. Whiskers indicate 95% confidence intervals. A significant inverse correlation was observed showing r=−0.42, p=0.0149 at n=41 candidates for 99 MSI colorectal cancers with wild type *B2M*, with a conservative estimate of predicted epitope fidelity of p_binding_=50%, indicating that high epitope likelihood was related to lower mutation frequency.

### Immunoselection during MSI carcinogenesis

In order to systematically evaluate whether these observations may result from immunoediting, i.e. counterselection of emerging cancer cell clones that harbor highly immunogenic cMS mutations (high GELS *neo*antigens), we analyzed potential differences between the observed and expected distribution of cMS mutations. We observed a significant inverse correlation between GELS and mutation frequency with Pearson’s *r* = −0.45, *p* = 0.0078 at *n* = 41 cMS for endometrial tumors and *r* = −0.42, *p* = 0.0149 for colon tumors, with a conservative estimate of predicted HLA binding probability of *p*_*binding*_ = 50%, indicating that a high GELS was related to lower mutation frequency (Fig. 4C). The correlation remained significant even at the lowest epitope fidelity levels of *p*_*binding*_ = 10%, with *p* = 0.0145 for endometrial and *p* = 0.0031 for colon cancers respectively.

The observation suggests that emerging tumor cell clones with highly immunogenic neoantigens are counterselected (Fig. 5), showing for the first time that immunoediting leaves its traces in *neo*antigen/cMS mutation patterns in MSI cancers (*35–38*). Interestingly, the significant inverse correlation was only detected among *B2M*-wild type tumors. *B2M*-mutant tumors, in which immune selection on the basis of HLA class I antigen presentation should not apply, only a trend was observed (Fig. 4D), which possibly reflects effects of immune surveillance prior to *B2M* mutation (see Data S4 for detailed test parameters).

**Fig. 5.**
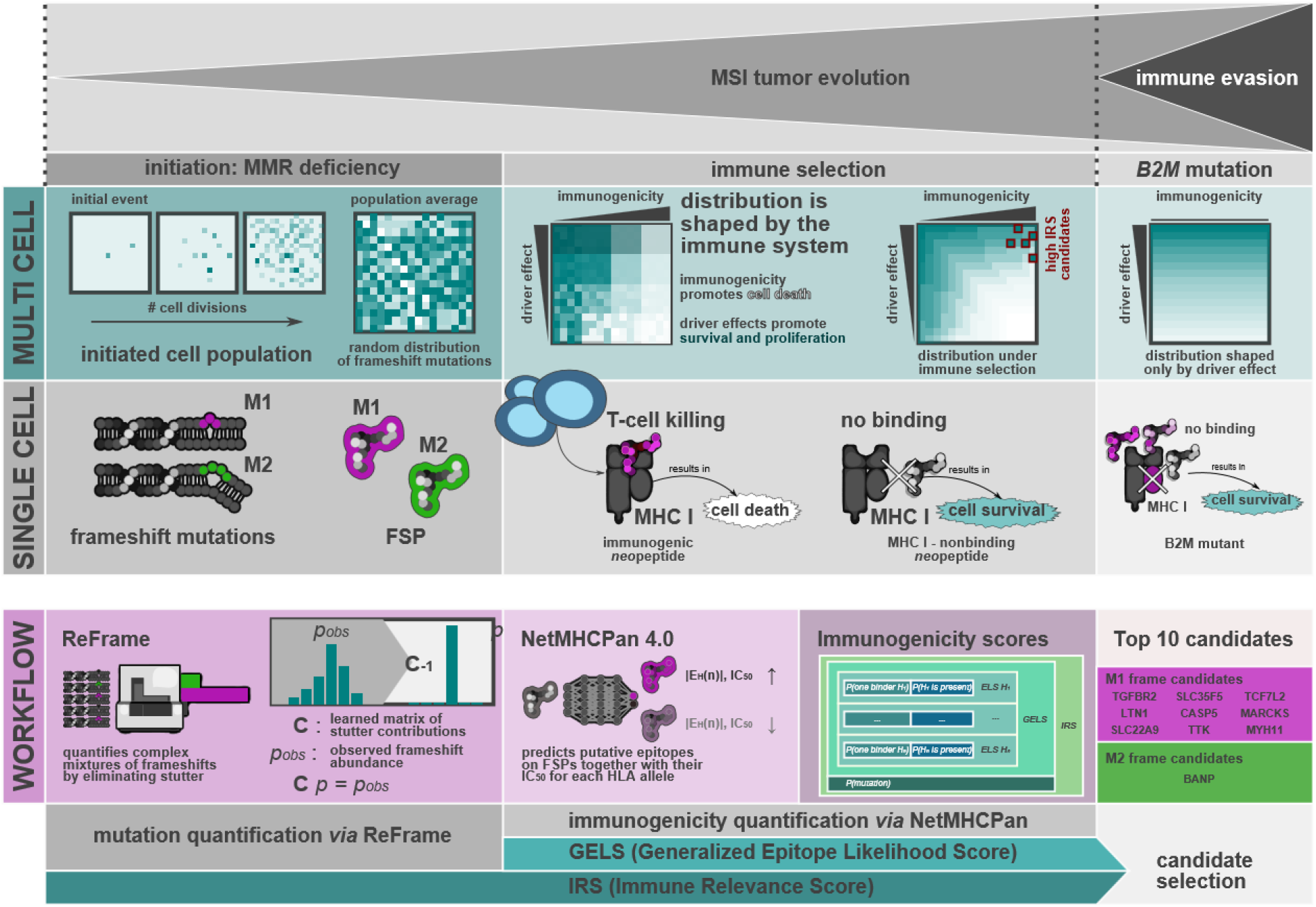
Implications of immune selection during tumor evolution in MSI cancers. **(MULTI CELL)** Inactivation of the MMR system results in the accumulation of a high number of somatic cMS mutations during cell division. These cMS mutation events depend on the likelihood of polymerase slippage at the microsatellite loci, i.e. on microsatellite length, but are random with regard to the functional consequences of the mutations, which results in a random distribution of cMS mutations in the initiated cell population. During progression, driver mutations promoting cell survival and proliferation are favorable, while highly immunogenic mutations are disavowable due to immune supervision. As such, the distribution of cMS mutations across a cell population is shaped by both driver effects and immune supervision. Abrogation of cellular antigen presentation, i.e. due to *B2M* mutation-induced loss of HLA class I stability, the immunogenicity of *neo*antigens resulting from cMS mutations is expected to become irrelevant for the selection of cell clones. Therefore, the distribution of cMS mutations is no longer shaped by the immune system and depends only on driver effects. (**SINGLE CELL**) Insertions and deletions due to polymerase slippage in cMS result in two equivalence classes of frameshift neopeptides with M1 or M2 frameshifts. Survival of a given cell with cMS mutations then depends on the binding behavior of these neopeptides to the cell’s HLA class I complexes. If *neo*antigens contain HLA binding peptides, they can be recognized as foreign by T cells, resulting in the possibility of T cell-mediated induction of cell death. In contrast, neoantigens not containing HLA binding peptides are neutral and do not impair cell survival. Destabilization of HLA class I by *B2M* mutation leads to a general lack of peptide-containing HLA class I complexes on the cell surface, theoretically corresponding to a complete lack of HLA class I binders. (**WORKFLOW**) Distribution of cMS mutations across tumor samples was quantified using ReFrame, which performs deconvolution on observed frameshift sequence abundances including stutter contributions to recover the true abundance of each frameshift sequence. NetMHCPan 4.0 predicts the IC50 of putative epitopes for all cMS-derived FSPs, identifying potential highly immunogenic FSPs by their number of predicted low-IC50 epitopes. This information is composed into a hierarchy of immunogenicity scores (ELS, GELS, IRS) combining multiple probabilities of HLA class I binding, presence of correct HLA types and presence of cMS mutations. The top 10 IRS FSPs are picked as possible candidates for vaccination.

We ruled out a potential influence of cMS length, a well-known factor influencing the likelihood of indel mutations on the observed mutation frequency (Fig. S54), (*23, 27, 39*) further supporting the concept of immunosurveillance-induced negative selection.

Despite the statistically significant negative correlation between GELS and mutation frequency, we also observed some outliers (Fig. 4C). We hypothesize that these outliers may reflect distinct effects that potentially influence the probability of a certain cell clone harboring a defined mutation to survive and thrive during tumor evolution. In addition to potential enhancement of immunogenicity, cMS mutations in tumor suppressor genes are predicted to lead to a growth advantage, at least in cancer or pre-cancer cell clones not directly under attack of the immune system. Such cMS candidates with high GELS and mutation frequencies should be of great relevance for the interaction between the immune system and MMR-deficient tumor cells. The presence of a *neo*antigen-inducing mutation is a prerequisite for presentation of corresponding *neo*epitopes that can be recognized by the host’s immune system. To simultaneously account for mutation frequency and GELS as factors influencing the likelihood of the *neo*antigen being presented to the immune system, we defined an “immune relevance score” (IRS), which combines GELS with the mutation frequency in tumors computed via ReFrame (see Materials and Methods section “Computation of immunological scores“).

The M1 FSP *neo*antigen derived from TGFBR2, the first described cMS driver mutation in MSI cancer and also the first ever FSP *neo*antigen characterized for its immunological properties in MSI cancer in pioneering studies (*18, 40, 41*), displays the highest IRS (28.57%). In addition to this well-characterized FSP *neo*antigen, our study uncovered various novel candidates with predicted importance for the immune biology of MMR-deficient cancers. The candidates LTN1, SLC22A9, SLC35F5, CASP5, TTK, TCF7L2, MYH11, MARCKS (all M1) and BANP (M2) all displayed an IRS above 10% (Fig. 4C, Data S3). The spatial distribution of predicted HLA-binding peptides within these high-IRS FSP *neo*antigens is visualized in Fig. S6. Interestingly, candidate genes with a possible tumor suppressor function were common among the high-IRS genes: *CASP5* (apoptosis induction; IRS: 17.15%), *TTK* (maintenance of chromosomal stability; IRS: 12.38%), *TCF7L2* (beta-catenin signaling; IRS: 11.32%), *MYH11* (cell structure and proliferation; IRS: 11.11%) and *BANP* (migration and invasiveness; IRS: 10.73%) were all previously reported in the literature (*42–50*). This observation may suggest that highly immunogenic *neo*antigens are ‘tolerated’ preferentially if the cells gain a compensatory survival advantage from the mutation by switching off a tumor-suppressive pathway, supporting their role of propelling MSI tumor evolution (Fig. 5).

## Discussion

MMR-deficient tumors, due to their well-defined mechanism of genomic instability, represent an ideal tumor type to study the evolution of solid cancer development and the role of the immune system during this process. By analyzing a broad spectrum of cMS-encompassing genes that are susceptible to mutation in MMR-deficient cells, we were able to identify recurrent mutations and *neo*antigens, and to provide first evidence for immunoediting during MSI cancer development.

The results of our study (Fig. 5) demonstrate that, in contrast to *neo*antigens in many other cancer types, which are typically differing between tumors or even occur as ‘private’ mutational *neo*antigens, MMR-deficient cancers share a large pool of FSP *neo*antigens. Thereby, most of the alterations are of the M1 type, resulting from one-basepair deletions (*m1*), with several candidates displaying a high likelihood of immunogenicity. This observation points towards a common evolutionary pathway of MSI tumorigenesis. The apparent dominance of *m1* mutations emphasizes that MMR-deficient cancers not only share similar sets of genes inactivated by MMR deficiency-induced mutations, but also precisely the same FSP *neo*antigens resulting from these mutations, allowing the definition of a shared *neo*antigen set for MMR-deficient cancers.

Using NetMHCpan 4.0, we identified a plethora of potential MHC binding peptides in FSP *neo*antigens. This number may even increase when using looser prediction thresholds, as recommended in a recent study evaluating the performance of MHC ligand prediction tools (*51*). Although many FSP *neo*antigens do not encompass such peptides for any of the common HLA types, our calculations demonstrate that the vast majority of MSI cancers are predicted to generate one or more *neo*antigens potentially recognizable by the host’s immune system. This hypothesis is supported by the observation of common FSP *neo*antigen-specific T cell responses in patients with MSI cancer and Lynch syndrome mutation carriers (*8*). As demonstrated by previous studies, even very low-affinity peptides may encompass relevant epitopes (*52, 53*). Moreover, several of the FSP *neo*antigens derived from common cMS mutation encompass “hot spot sequences” for which multiple HLA-binding peptides have been predicted (indicated by dark colors in Fig. 3), suggesting that these might be of increased interest for further evaluation (*52, 53*).

Our study has the following limitations. The list of *neo*antigens analyzed with ReFrame is not exhaustive, as additional frameshift mutations resulting from shorter, less frequently mutated cMS can occur in MSI cancers. In addition, we can only propose an atlas of predicted potential *neo*epitopes in MSI cancers. Although previous studies evaluated a few of the predicted candidates (*18, 54*), supporting the general validity of the *in silico* predictions, functional validation of individual predicted epitopes will be required to demonstrate that they can in fact be processed by tumor cells and recognized by immune cells.

By combining quantitative cMS mutation analysis with a *neo*antigen-specific immune score that accounts for the prevalence of the epitope-binding HLA molecules in the population, we for the first time are able to provide evidence that the cMS mutation patterns in MSI cancers show signs of immune selection: Candidates that encompass immunogenic epitopes predicted to bind to common HLA types tend to occur less frequently in manifest MSI cancers. This observation supports the concept that immune surveillance is a major force shaping the natural course of MMR-deficient cancer development (*4, 25, 26, 37, 55*). Depletion of expressed neoantigens, similar to what our data suggest, has recently been reported in lung cancer (*56*).

Other studies failed to detect evidence for negative selection of immunogenic, neoantigen-inducing mutations in cancer and thereby immunoediting (*57, 58*). This discrepancy may in part be related to the fact that our approach specifically compares individual cMS mutations based on their immunological consequences, accounting not only for the presence of predicted epitope sequences, but also for the population frequency of the respective HLA type, to which the predicted epitope is supposed to bind. In addition, the detectability of specific counterselection events is supported by three specific features of MMR deficiency: first, MMR-deficient cancers in contrast to other tumors share precisely the same mutations, because the location of a cMS within a gene determines its susceptibility for indel mutations in MMR-deficient cells; second, MMR-deficient cancers due to the dramatically elevated rate of somatic mutations per cell division are expected to harbor a significantly higher proportion of MMR deficiency-induced mutations compared to age-related mutations that have occurred prior to tumor initiation, thus enhancing the “visibility” of negative selection events; third, counterselection against FSP *neo*antigens may be particularly pronounced, as MMR deficiency-induced mutations often lead to generation of long *neo*antigens with potentially multiple epitopes, against which no central immune tolerance exists (*59*).

The observation of immunoediting during the development of MMR-deficient cancers also implies that a person’s HLA genotype should have a significant influence on the immune environment during MSI tumor evolution. Given the existence of immune-relevant FSP *neo*antigens that may be bound only by a certain type of HLA molecules, it is reasonable to assume that HLA genotype may be a modifier of cancer risk. This may also explain possible variations of Lynch syndrome penetrance or different rates of MMR deficiency previously suspected between distinct populations (*60*). Future studies on the natural course of Lynch syndrome should account for this factor.

The shared *neo*antigen landscape encourages cancer-preventive vaccines against MSI cancers, particularly in the setting of Lynch syndrome. If we are able to enhance the abundance of T cells recognizing FSP *neo*antigens by an FSP *neo*antigen vaccine, we may shift the balance towards elimination of emerging cancer cells, thereby reducing the likelihood of escape variants leading to outgrowth of clinically manifest tumors. The safety and immunological efficacy of such an FSP *neo*antigen-based vaccine has already been demonstrated in a first clinical phase I/IIa trial (https://clinicaltrials.gov/show/NCT01461148). If the immune system can be specifically sensitized towards FSP *neo*antigens resulting from driver mutations which inactivate tumor suppressor genes, such as the ones we evaluated in this study, tumor evolution should be influenced in a way that outgrowth of ‘dangerous’ MSI cancer cell clones should become significantly less likely.

In conclusion, mutational landscapes in MSI cancers suggest negative selection of mutations that give rise to highly immunogenic FSP *neo*antigens. This supports the validity of the immunoediting concept in non-viral human tumors. *Neo*antigen-based vaccination approaches for the prevention of MMR-deficient cancers should account for the natural immune surveillance during their development and focus on strengthening the host’s immune response against *neo*antigens that are related to essential driver mutation events.

## Materials and Methods

### Tumor specimens

Formalin-fixed, paraffin-embedded (FFPE) archival tissue blocks were collected from 139 MSI colorectal carcinomas and 14 MSI endometrial carcinomas. Pseudonymized clinical data of each tumor patient is summarized in Table S1. Tumors were obtained from the Department of Applied Tumor Biology, University Hospital Heidelberg in frame of the German HNPCC Consortium, the Finnish Lynch syndrome registry, and Leiden University Medical Center. The study was approved by the Institutional Ethics Committee, University Hospital Heidelberg. Informed consent was obtained from all patients.

### Tissue workup and DNA isolation

FFPE tumor sections (5 μm) were deparaffinized and stained with hematoxylin and eosin according to standard protocols. DNA was isolated from tissue sections after separate microdissection of normal and tumor tissue. Only samples with a tumor cell content of more than 80% were used for the analysis. Genomic DNA was isolated using the Qiagen DNeasy Tissue Kit (Cat.No. 69506, Qiagen, Hilden, Germany) according to the manufacturer’s instructions.

### MSI analysis

The tumors were characterized for their MSI status using the NCI/ICG-HNPCC five microsatellite marker panel supplemented with additional mononucleotide markers BAT40 and CAT25 (*61*). Tumors displaying instability in more than 30% of the analyzed markers were classified as MSI.

### Analysis of frameshift mutations in coding microsatellites (cMS)

In order to amplify the coding microsatellite loci, primers were either obtained from the Seltarbase (http://www.seltarbase.org) (*27*) or designed using primer3 software (Primer3web version 4.0.0, http://primer3.ut.ee/), with one primer of the primer set carrying a 5’ fluorescent (FITC) label. Primer were designed to generate amplicons in range between 100 and 150 nucleotides for robust PCR amplification (Table S7). PCR was performed in a total volume of 5 μl containing 0.5 μl 10x reaction buffer (Invitrogen, Karlsruhe, Germany), 1.5 mM MgCl_2_, 200 mM dNTP mix, 0.3 mM of each primer, 0.1 U Taq DNA polymerase (Invitrogen), and 10 ng of genomic DNA, using the following protocol: initial denaturation at 94°C for 5min; 36 cycles of denaturation at 94°C for 30s, annealing at 58°C for 45s and primer extension at 72°C for 1min; final extension step at 72°C for 7min. PCR fragments were separated on an ABI3130*xl* genetic analyzer (Applied Biosystems, Darmstadt, Germany). Generated raw data were analyzed using GeneMapper™ Software version 4.0 (ThermoFisher, Waltham, USA). Peak height profiles were extracted and processed using ReFrame based on R version 3.4.3. The R script is available as Supplementary Material 1.

### Microsatellite allele distributions analyzed using Regression-based Frameshift quantification (ReFrame)

In general, PCR amplification of microsatellite loci generates fragments that can vary in length, either due to indel mutations in MMR-deficient cells or due to polymerase slippage during amplification (stutter band artifacts). These two phenomena cause overlays of peak patterns and hamper data interpretation. We developed a ReFrame, a REgression-based FRAMEshift quantification algorithm, to allow quantitative analysis of microsatellite mutations by removing stutter band artifacts.

We obtained main-peak fractions as a function of microsatellite length, to which a logistic function, in the following referred to as *p*(*L*) was then fitted. For each microsatellite in question, an effective length was computed using that fit. We then determined stutter fractions for each gene, by calculating the ratios of additional fragments occurring at each microsatellite locus in MMR-proficient control samples (n = 20) to establish baseline reference values *p*_*ref*_. For each cMS, we computed the expected relative contributions of each insertion/deletion in the range of *Δ* = −4 deletion to *Δ* = +4 insertion to each band in the data as:

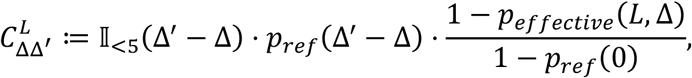

where we defined

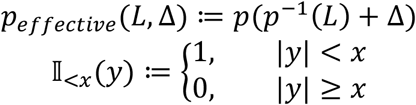

We used these relative contributions to set up a linear system for the true peak size without stutter contributions (Fig. S6) by requiring

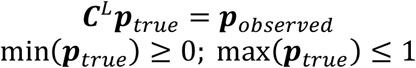

where ***p***_*observed*_ and ***p***_*true*_ are the observed and true peak sizes respectively. Resulting allele profiles were imported into a database for further analysis.

Validation of ReFrame was performed in three steps: First, DNA of colonic normal tissue was used to determine baseline deviations of the method in negative controls (Fig. S6c). Additionally, microsatellite-stable cell line DNA (HT29) was used as a control. Finally, two cell line DNAs with differing mutation states (HT29 displaying wild type peak pattern, LS180, displaying a mutant peak pattern) were mixed in 10%-steps and expected allele distributions were compared to the ReFrame results (Fig. S6d).

### Code availability

The source code of all used algorithms can be accessed on https://github.com/atb-data/neoantigen-landscape-msi

### Selection of coding microsatellites and frameshift peptide sequences

For HLA class I binding prediction, 524 FSP *neo*antigen sequences from 262 mononucleotide changes were retrieved from the Selective Targets in Human MSI-H Tumorigenesis Database (Seltarbase, http://www.seltarbase.org) (*27*). All cMS with a length of at least eight bases were included. In particular cases other cMS representing putative driver genes, as well as genes which give rise to FSPs with predicted high-affinity binding epitopes according to the literature were also added to the study. In order to also assess potential epitopes located at the junction between N-terminal wild type and C-terminal mutant peptide sequences, the tested peptide sequences all comprised 8 wild type amino acids directly located upstream of the FSP *neo*antigen sequence to encompass possible fusion epitopes. The whole list of used FSP *neo*antigens is depicted in Table S5.

### HLA binding predictions

For HLA binding prediction, the *neo*antigen sequences derived from each the M1 and M2 mutated alleles were analyzed for the presence of binders using the publicly available prediction tool NetMHCpan 4.0 (www.cbs.dtu.dk/services/NetMHCpan/) (*9*), whose performance has been evaluated to be one of the best of the available tools (*51*). As *m1*-induced and *p2*-induced M1 *neo*antigens (akin to *m2*-induced and *p1*-induced M2 *neo*antigens) are identical, except for one additional amino acid at the transition between wild type and *neo*-sequence, we only used M1/*m1* and M2/*m2* neoantigens for HLA binding prediction.

Predicted epitopes were subdivided into three classes based on commonly used thresholds. While the first class included epitopes with a predicted affinity of IC_50_ below 50 nM, referred to as high-affinity binders, the second class included all predicted binders below 500 nM (low-affinity binders). The last class was containing all putative epitopes with lower than 5000 nM affinity (very low-affinity binders). All potential HLA binders with an affinity higher than 5000 nM were discarded. The peptide length of interest was set to 8mer to 14mer peptides. A preselection of HLA supertype representatives including HLA-A*01:01, HLA-A*02:01, HLA-A*03:01, HLA-A*24:02, HLA-A*26:01, HLA-B*07:02, HLA-B*08:01, HLA-B*27:05, HLA-B*39:01, HLA-B*40:01, HLA-B*58:01 and HLA-B*15:01 was chosen based on previous recommendations (*31, 32*). A list of all chosen cMS and FSP sequences were submitted to a Python driver script operating NetMHCpan 4.0 (*9*) to predict putative HLA binding peptides. The prediction results were processed using a Python script applying the above-mentioned IC50 thresholds to all predicted peptides, yielding three datasets of peptides with potential very low, low and high HLA binding affinity. The resulting datasets were then used to generate figures visualizing the predicted epitopes using matplotlib (*62*). To that end, predicted epitopes were counted and mapped for each HLA type, *neo*antigen candidate and the respective epitope class (high-, low- or very low-affinity binder). The results of that analysis were used to generate heatmaps per candidate and HLA type using another Python script (see Suppl. Material 1 for all scripts).

### Selection of HLA allele frequency data

HLA allele frequency data sets were selected from the Allele Frequency Net Database (*34*) by taking the largest datasets of each ethnicity with at least 10000 data points and sufficient resolution in HLA alleles. These were further processed together with epitope and mutation data to compute the immunological scores.

### Computation of immunological scores

For all candidate FSP *neo*antigen, measures of probable immunological relevance were computed based on the above described predicted IC_50_ values and mutation frequencies. A hierarchy of probabilities for the given candidates to produce immune reactions were computed, those being an epitope likelihood score (ELS) per HLA type, a generalized epitope likelihood score (GELS) comprising all HLAs under consideration, as well as an immunological relevance score (IRS). The ELS was defined to describe the probability of a given *neo*antigen to be effective across a population, relative to a single HLA:

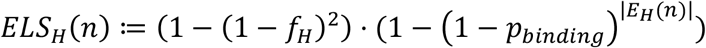

where *H* ∈ *HLA supertypes* is a given HLA, *n* ∈ *cMS* is a given FSP *neo*antigen, *f*_*H*_ the allele frequency of a given HLA allele, *p*_*binding*_ the probability, that a given predicted epitope is actually bound, that is the true positive rate of the prediction algorithm, and *E*_*H*_(*n*) the set of all epitopes predicted for a given HLA and *neo*antigen. Taken together, *ELS*_*H*_ constitutes the probability of a given candidate *n* having at least one true binding epitope for an HLA *H* and a random person from a given population having at least one allele of *H*.

Consequently, the GELS gives the probability of a candidate *n* having at least one binding epitope among all HLAs, for which the given HLA is also present in a randomly selected individual:

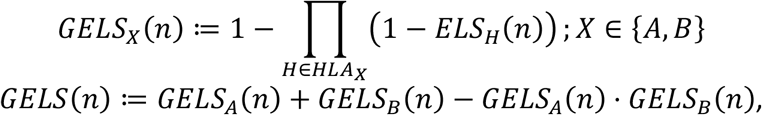

where *HLA*_*X*_ is the set of HLA types considered for locus *X*.

Finally, the IRS gives the joint probability of a given FSP and its underlying cMS mutation being present in an individual and at least one predicted binder existing for an HLA present in that individual, assuming independence between the presence of HLA alleles and present FSPs:

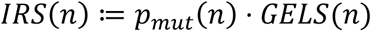

ELS and GELS were computed for all candidate FSPs and HLAs considered using Python on the three output classes of epitope prediction, where binding probabilities *p*_*binding*_ were incremented from 0% to 90% in steps of 10%. HLA allele frequencies were obtained from the Allele Frequency Net Database (*34*). Immunological relevance scores were computed for all candidates with available mutation frequency data.

### Cluster analysis of mutation patterns

Frameshift mutation abundances (m4 to p4) for each gene and tumor sample were filtered for missing data. For all subsequent clustering experiments, missing values were replaced by the dataset mean. Abundances of frameshift mutations were summarized by their respective reading frame (M2, M1, wt), providing the features used for all subsequent analyses. Resulting features were grouped by tumor sample and candidate cMS respectively. Hierarchical clustering using Ward’s minimum variance linkage (*63*) was performed for both feature-sets grouped by cMS and tumors for all tumor samples considered, as well as for cMS features considering only *B2M* wildtype and mutated tumors respectively. Three clusters of candidate cMS were extracted from hierarchical clustering both for features considering all tumor samples and features considering *B2M* wildtype tumors only.

## Glossary

ELS: Epitope likelihood score
GELS: General epitope likelihood score
IRS: Immune relevance score, based on the mutation frequency (ReFrame) and the GELS
M1: Reading frame resulting from the deletion of one nucleotide or insertions of two nucleotides
M2: Reading frame resulting from the deletions of two nucleotides or insertion of one nucleotide
*m1*, *m2*, *m3*, etc.: Minus one, two, three base pair deletions
*p1*, *p2*, *p3*, etc.: Plus one, two, three base pair insertions
ReFrame: REgression-based FRAMEshift quantification

## Abbreviations

B2M: Beta2-microglobulin
cMS: Coding microsatellites
CRC: Colorectal cancer
MMR: Mismatch repair
EC: Endometrium cancer
FSP *neo*antigens: Frameshift peptide *neo*antigens
HLA: Human leukocyte antigen
ICB: Immune checkpoint blockade
MSI: Microsatellite instability, microsatellite-unstable
NGS: Next generation sequencing

## Acknowledgements

The excellent technical assistance of Nina Nelius, Petra Hoefler, and Beate Kuchenbuch is gratefully acknowledged.

## Author contributions

Study conception: AB, MJP, MJ, MKD, MK

Molecular analysis: AB, MJP, EP, MD, FS, SK, AA

Data analysis: AB, MJP, MJ, SH, EP, MD, FS, SK, AA, MSK, DH, MB, VH, JK, ABe, ABR, MK

Data interpretation: AB, MJP, MJ, SH, MSK, DH, JG, MB, SS, HB, VH, JK, ABe, ABR, MKD, MK

Manuscript writing: AB, MJP, MJ, SH, JG, MB, MKD, MK

Providing tissue specimens: SS, HB, TS, JPM, STB, MN, MKD, MK

Revision and final approval of the manuscript: All authors

## Data and materials availability

The source code of all used algorithms with the corresponding data Data S1 to S5 can be found on https://github.com/atb-data/neoantigen-landscape-msi

## Supplementary Materials

Figs. S1 to S6

Tables S1 to S5

Captions for Data S1 to S7

