## Supplementary Material for "The shared *neo*antigen landscape of MSI cancers reflects immunoediting during tumor evolution"

$ shared first authorship

Affiliations:
1 Department of Applied Tumor Biology, Institute of Pathology, University of Heidelberg, Heidelberg, Germany.
2 Collaboration Unit Applied Tumor Biology, German Cancer Research Center (DKFZ), Heidelberg, Germany.
3 Molecular Medicine Partnership Unit (MMPU), Heidelberg University Hospital and EMBL Heidelberg, Germany.
4 Engineering Mathematics and Computing Lab (EMCL), Interdisciplinary Center for Scientific Computing (IWR), Heidelberg University, Heidelberg, Germany
5 Immunotherapy and Immunoprevention, German Cancer Research Center (DKFZ), Heidelberg, Germany
6 Molecular Vaccine Design, German Center for Infection Research (DZIF), partner site Heidelberg, Heidelberg, Germany
7 Faculty of Biosciences, Heidelberg University, Heidelberg, Germany
8 Department of Obstetrics and Gynecology, University Hospital Heidelberg, Heidelberg, Germany.
9 Institute of Pathology, University Hospital Leipzig, Leipzig, Germany.
10 Department of Gastrointestinal Surgery, Helsinki University Hospital and University of Helsinki, Helsinki, Finland
11 Department of Education and Research, Central Finland Central Hospital, Jyväskylä, Finland, and Sports and Health Sciences, University of Jyväskylä, Jyväskylä, Finland
12 Department of Clinical Genetics, Leiden University Medical Center, Leiden, The Netherlands
13 Division of Biostatistics, German Cancer Research Center (DKFZ), Heidelberg, Germany

* Corresponding author

**This PDF file includes:**

Figs. S1 to S6
Tables S1 to S6
Captions for Data S1 to S5

**Other Supplementary Materials for this manuscript include the following:**

Data S1 to S5, Data S1 - cMS peptide sequences, Data S2 - Predicted HLA binding peptides per candidate and HLA, Data S3 - GELS and IRS in various populations, Data S4 - ReFrame and epitope prediction scripts, Data S5 - Plots depicting predicted HLA binding peptides for all FSP neoantigens.

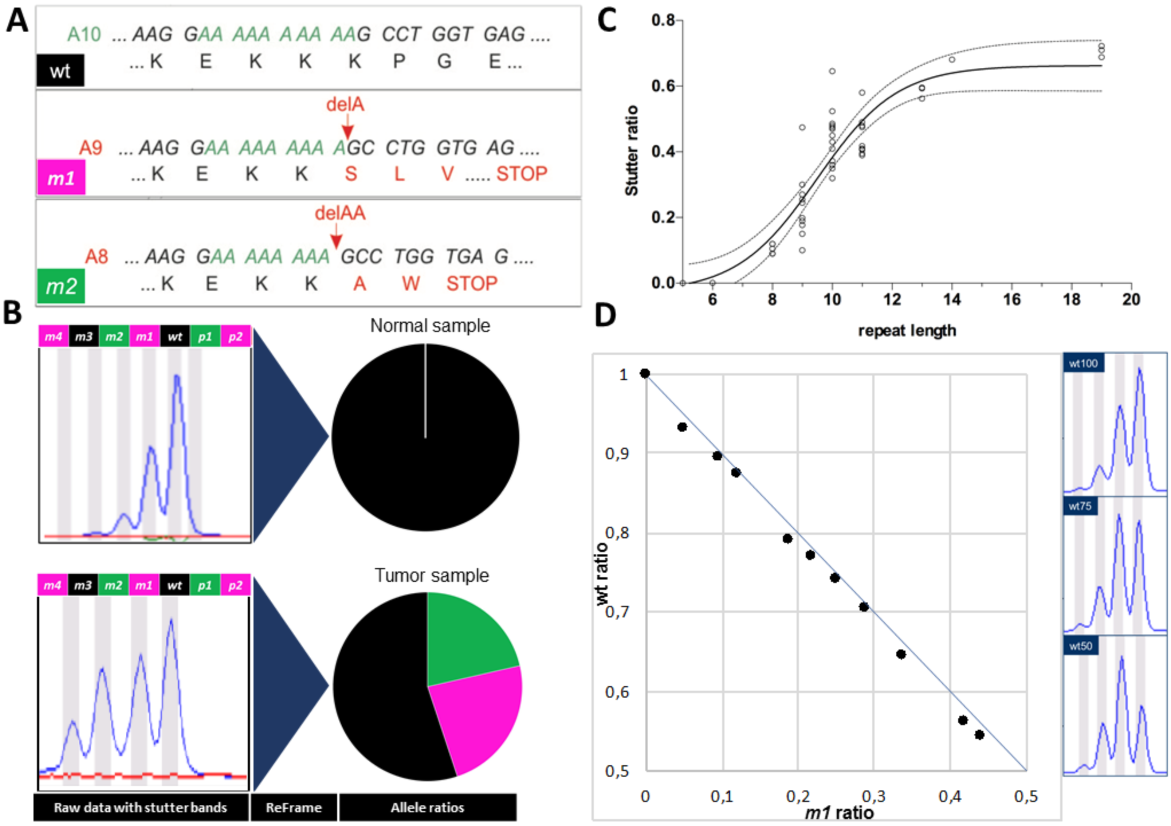


**Fig. S1. Concept and validation of REgression-based FRAMEshift quantification (ReFrame) algorithm**. (A) A10 cMS of the *TGFBR2* gene, including amino acid translation of wild type, *m1* (red) and *m2* (green) alleles. (B) Peak profile obtained from colonic normal tissue with a wild type *TGFBR2* A10 allele (upper panel). For comparison, the peak profile of matching tumor tissue, carrying *m1* and *m2* mutations is shown (lower panel). Pie charts illustrate allele frequency after removal of stutter bands by ReFrame. (C) Stutter ratios of raw data normal tissues are plotted against cMS repeat length. (D) MSI cell lines with known mutation status of a defined cMS (NDUFC2, LS180: m1/wt, HT29: wt) were mixed to compare experimental ReFrame results (black dots) with theoretically expected values (blue line). Three representative peak patterns (raw data prior to stutter band removal) are shown the right panel.

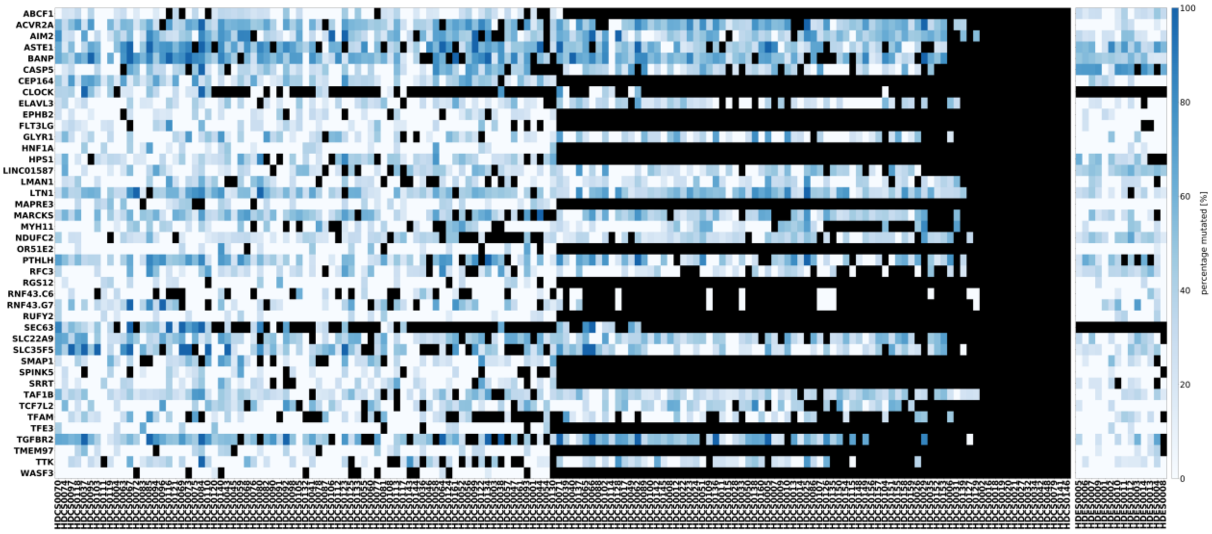


**Fig. S2. Mutation frequencies of coding microsatellites (cMS) in MSI CRC and EC.** The relative frequency of mutant alleles analyzed by the qMSI algorithm is shown for 41 cMS (rows) in all CRC and EC tumor samples (columns). using ReFrame. cMS were sorted alphabetically. Dark blue represents high mutation frequency, whereas pale blue represents low mutation frequency. Black boxes indicate missing data points. cMS were analyzed for both CRC and EC and are depicted separately for each tumor type (left panel: CRC, right panel: EC).

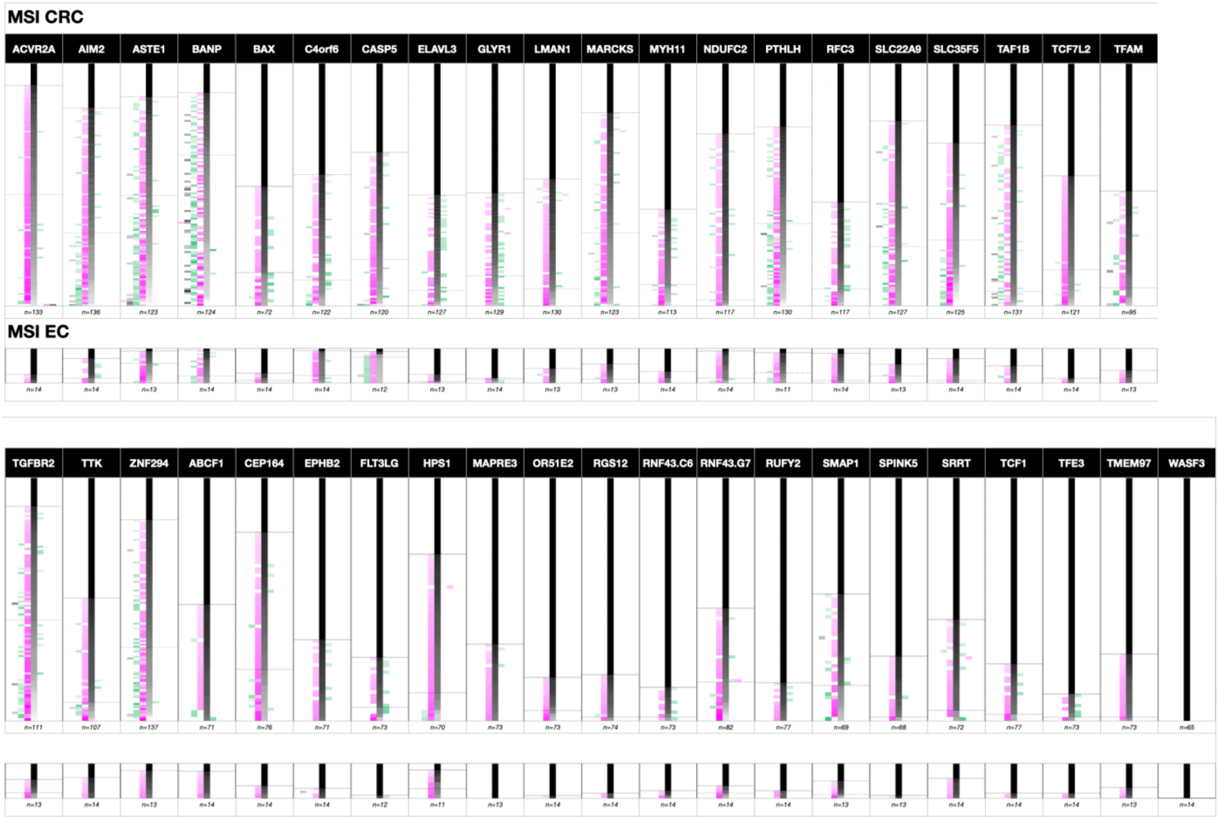


**Fig. S3. Mutational pattern distribution in cMS based on ReFrame analysis.** The detailed mutational patterns of all 41 analyzed cMS are represented with their respective frequency of mutation for all possible resulting frameshift mutations in MSI CRC and EC. Each row constitutes one analyzed tumor sample with its related allele ratios. The number of samples analyzed for a certain candidate is indicated below for each candidate. Since wt, m3 and p3 mutations do not result in translational frameshifts, they are shown in black. In contrast, m1, m4 and p2 mutations (red) and m2, p1, p4 mutations (green) are either resulting in a frameshift peptide arising from a one base pair or two base pair deletion reading frame, respectively. The column intensities represent calculated ratios from white (0%) to the respective color of the column (100%). All samples are sorted according to their wild-type proportion. The annotated solid lines show the end of the non-mutated tumor samples while the dotted lines mark the beginning of tumors that are mutated in more than 50 % of cases, associated with biallelic hits within the respective sample. Some candidates which were analyzed in CRC were not analyzed for EC (white spaces).

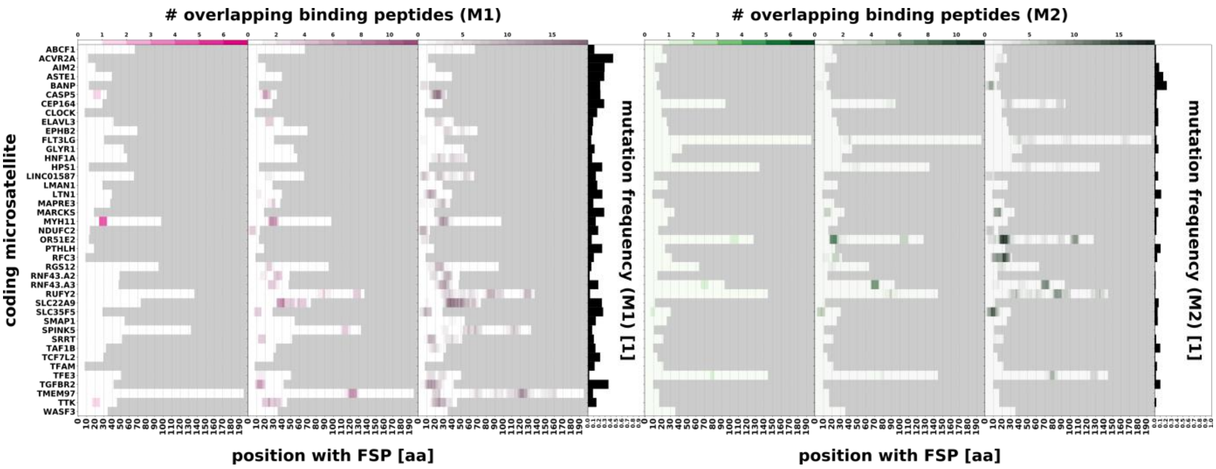


**Fig. S4. Epitope predictions in HLA-A*02:01.** The figures display the predicted epitopes in HLA-A*02:01, the most frequent HLA-type in the western world, for the m1 and m2 FSPs of all 41 cMS candidates over the length of the respective peptide. High-affinity ($IC_{50}$ < 50 nM), low-affinity ($IC_{50}$ < 500 nM), and very low-affinity epitopes ($IC_{50}$ < 5000 nM), are shown for the m1 FSPs (left, red) and the m2 FSPs (right, green). All candidates are sorted in alphabetical order with their respective frequency of mutation according to the results of the ReFrame algorithm. The grey field is highlighting the ends of the respective FSPs.

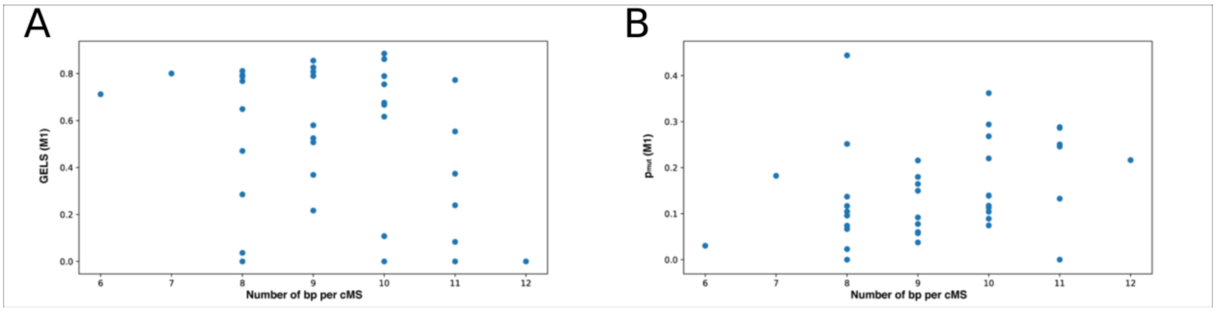


**Fig. S5. GELS and mutation frequency ($p_{mut}$) in dependency on cMS length.** (**A**), All cMS which were analyzed using ReFrame, thus having a calculated IRS, are depicted according to their number of base pairs on the x-axis and their GELS on the y-axis. The GELS is shown for the M1 frame only. (**B**), All cMS which were analyzed using ReFrame are depicted according to their number of base pairs and their mutational frequency on the y-axis. The mutational frequency is shown for the M1 frame only.

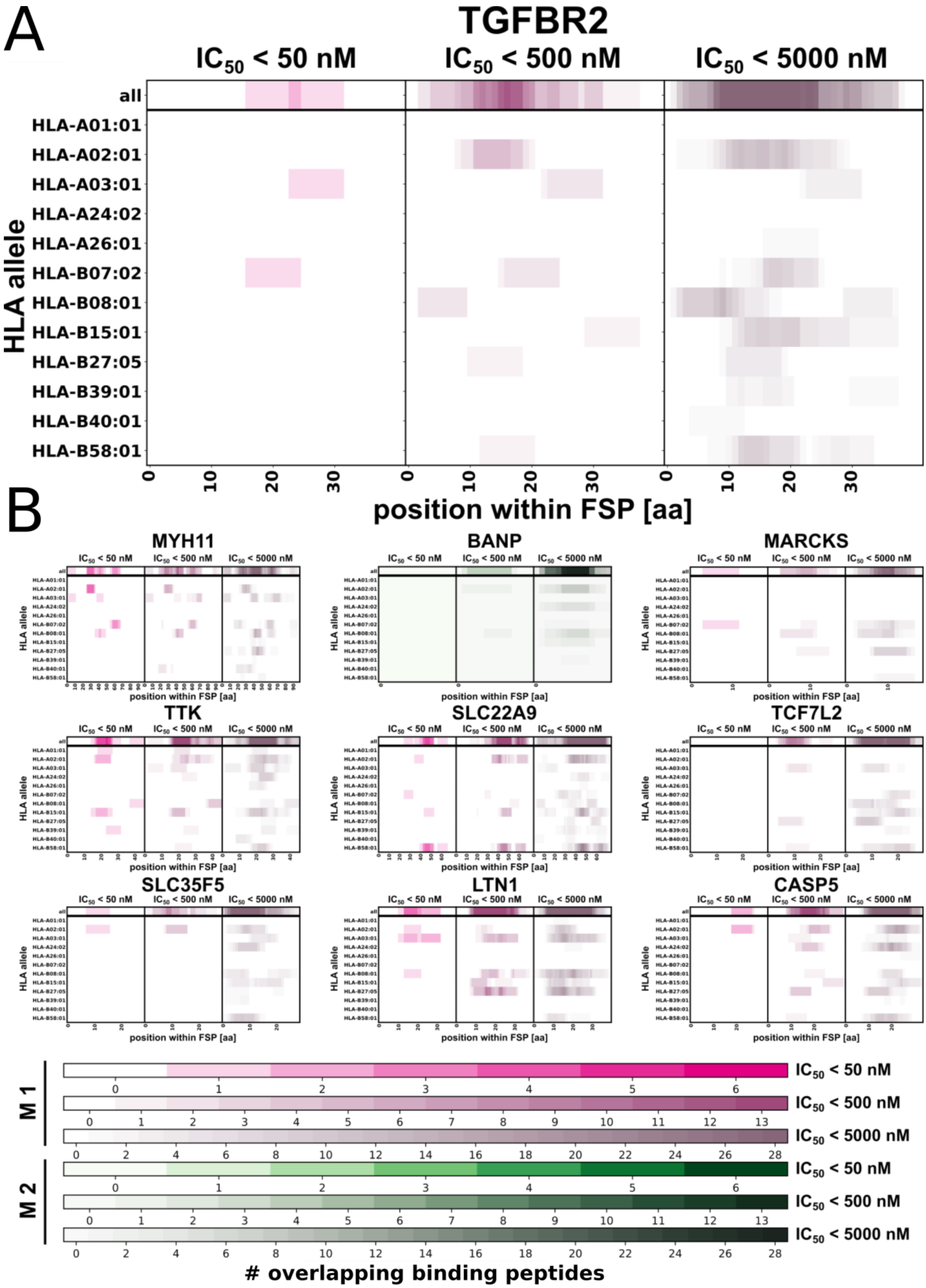


**Fig. S6. HLA binding affinity predictions for peptides derived from 10 cMS *neo*antigens.** (**A**) The epitope predictions for the M1 FSP *neo*antigen of TGFBR2 is shown for the HLA supertype representatives. The epitope prediction for the respective HLA type over the length of the neoantigen is shown in each column. Summarized hot spots of predicted HLA binding are shown at the top of each figure. For each candidate, the three different parts of the figure represent peptides with regard to their predicted HLA binding affinity (left – high, middle – low, right – very low). The position of the individual epitope is depicted on the x-axis, together with the length of the respective peptide. The color intensities are described below for the respective M1 and M2 FSP neoantigens. (**B**) HLA binding predictions for the additional cMS from the Top 10 IRS, including M1 FSP *neo*antigens LTN1, MARCKS, SLC22A9, SLC35F5, MYH11, TTK, TCF7L2, CASP5 and the BANP M2 FSP *neo*antigen are depicted for the HLA supertype representatives (see Data S6 and S7 for extensive information on all FSP *neo*antigens).

| gene name | cMS length (nucleotides) | aa change | DFCI MSI | Genentech MSI | TCGA MSI | Giannikis 2016 | Hause 2016* | Cortes-Ciriano 2017* | Kondelin 2017 (MiSeq93)* | Kondelin (MiSeq24)* | Kondelin (SANGER) | ReFrame* |
|---|---|---|---|---|---|---|---|---|---|---|---|---|
| BANP | 12 | p.L140fs | 0% | 0% | 0% | 0% | 0% | 0% | 0% | 0% | n.a. | 88% |
| ASTE1 | 11 | p.R657fs | 0% | 0% | 0% | 0% | 0% | 62% | 0% | 13% | 93% | 86% |
| LTN1 | 11 | p.N582fs | 0% | 0% | 0% | 0% | 59% | 47% | 0% | 13% | 89% | 83% |
| MARCKS | 11 | p.K155fs | 0% | 0% | 0% | 0% | 0% | 0% | 0% | 8% | n.a. | 80% |
| CEP164 | 11 | p.K70fs | 0% | 0% | 0% | 0% | 0% | 13% | 0% | 4% | n.a. | 78% |
| SLC22A9 | 11 | p.K335fs | 0% | 0% | 0% | 0% | 62% | 58% | 0% | 21% | n.a. | 76% |
| TAF1B | 11 | p.N66fs | 0% | 0% | 0% | 0% | 0% | 0% | 0% | 13% | n.a. | 75% |
| PTHLH | 11 | p.K186fs | 0% | 0% | 0% | 0% | 0% | 0% | 0% | 0% | n.a. | 74% |
| TGFBR2 | 10 | p.K128fs | 0% | 7% | 42% | 0% | 45% | 60% | 97% | 83% | 95% | 88% |
| AIM2 | 10 | p.K343fs | 0% | 0% | 26% | 0% | 27% | 60% | 0% | 8% | 78% | 82% |
| SLC35F5 | 10 | p.F248fs | 0% | 0% | 0% | 0% | 23% | 20% | 76% | 54% | n.a. | 67% |
| CASP5 | 10 | p.K68fs | 0% | 0% | 16% | 0% | 0% | 18% | 89% | 13% | 91% | 63% |
| LINC01587 | 10 | p.F24fs | 0% | 0% | 0% | 0% | 0% | 27% | 0% | 0% | n.a. | 54% |
| SMAP1 | 10 | p.K145fs | 0% | 0% | 0% | 0% | 0% | 60% | 0% | 21% | 92% | 52% |
| ABCF1 | 10 | p.K76fs | 0% | 0% | 0% | 0% | 0% | 0% | 0% | 33% | n.a. | 48% |
| TFAM | 10 | p.K147fs | 0% | 0% | 0% | 0% | 0% | 7% | 88% | 58% | n.a. | 47% |
| RFC3 | 10 | p.K82fs | 0% | 0% | 0% | 0% | 0% | 0% | 0% | 17% | n.a. | 43% |
| TMEM97 | 10 | p.K176fs | 0% | 0% | 0% | 0% | 0% | 18% | 0% | 17% | n.a. | 27% |
| SPINK5 | 10 | p.K823fs | 0% | 0% | 0% | 0% | 0% | 33% | 0% | 25% | n.a. | 26% |
| NDUFC2 | 9 | p.F69fs | 0% | 0% | 3% | 0% | 0% | 60% | 0% | 8% | n.a. | 71% |
| TCF7L2 | 9 | p.K462fs | 0% | 0% | 3% | 0% | 0% | 20% | 45% | 54% | n.a. | 54% |
| LMAN1 | 9 | p.K305fs | 0% | 0% | 0% | 0% | 51% | 33% | 49% | 38% | n.a. | 52% |
| TTK | 9 | p.G853fs | 0% | 0% | 0% | 1% | 0% | 24% | 47% | 58% | n.a. | 50% |
| ELAVL3 | 9 | p.G16fs | 0% | 0% | 0% | 0% | 0% | 0% | 0% | 0% | n.a. | 46% |
| EPHB2 | 9 | p.G1020fs | 0% | 0% | 0% | 0% | 0% | 33% | 0% | 0% | n.a. | 34% |
| FLT3LG | 9 | p.P118fs | 0% | 0% | 0% | 0% | 0% | 0% | 0% | 0% | n.a. | 26% |
| RGS12 | 9 | p.K1178fs | 0% | 0% | 0% | 0% | 34% | 22% | 0% | 4% | n.a. | 19% |
| RUFY2 | 9 | p.K131fs | 0% | 0% | 0% | 0% | 27% | 0% | 0% | 17% | n.a. | 16% |
| ACVR2A | 8 | p.K437fs | 21% | 0% | 47% | 4% | 0% | 84% | 88% | 92% | 89% | 91% |
| HPS1 | 8 | p.P325fs | 0% | 0% | 3% | 0% | 0% | 0% | 0% | 0% | n.a. | 69% |
| GLYR1 | 8 | p.G381fs | 0% | 0% | 0% | 0% | 0% | 0% | 0% | 33% | n.a. | 47% |
| SRRT | 8 | p.G106fs | 2% | 0% | 0% | 0% | 0% | 0% | 0% | 17% | n.a. | 42% |
| MYH11 | 8 | p.P1933fs | 0% | 0% | 0% | 0% | 10% | 0% | 0% | 0% | n.a. | 40% |
| MAPRE3 | 8 | p.P166fs | 0% | 0% | 3% | 0% | 25% | 0% | 0% | 29% | n.a. | 32% |
| HNF1A | 8 | p.P291fs | 2% | 0% | 0% | 0% | 0% | 0% | 0% | 8% | n.a. | 23% |
| OR51E2 | 8 | p.F156fs | 7% | 0% | 0% | 1% | 0% | 0% | 0% | 4% | n.a. | 18% |
| TFE3 | 8 | p.G482fs | 0% | 0% | 0% | 0% | 0% | 0% | 0% | 8% | n.a. | 11% |
| WASF3 | 8 | p.P307fs | 2% | 0% | 0% | 0% | 0% | 0% | 0% | 4% | n.a. | 0% |
| RNF43(G7) | 7 | p.G659fs | 24% | 7% | 3% | 5% | 0% | 36% | 0% | 8% | n.a. | 46% |
| RNF43(C6) | 6 | p.R117fs | 0% | 0% | 0% | 1% | 0% | 0% | 0% | 8% | n.a. | 14% |

**Table S1. Comparison of mutation frequencies obtained by ReFrame vs. other literature methods**. The mutational frequencies from the cohort of analyzed cMS candidates were compared to results from previous studies. Recent comprehensive next generation sequencing (NGS) based studies or online databases, where sufficient data were available, were included. Gene name, cMS repeat length and corresponding amino acid change are listed. The studies from Hause et al. (2016), Cortes-Ciriano et al. (2017) and Kondelin et al. (2017) as well as our ReFrame algorithm are marked with "*", as they show tailored approaches to enhance the detection of cMS mutations. In Kondelin et al. (2017), cMS that were not analyzed by Sanger sequencing are marked "n.a.".

| TU ID | MSI.Status | Tumor.type | Etiology (hereditary/sporadic) | age | sex | B2M.Seq | UICC | FIGO |
|---|---|---|---|---|---|---|---|---|
| HD001 | MSI | CRC | hereditary | 36 | female | mutated | 3 | NA |
| HD002 | MSI | CRC | hereditary | 71 | female | mutated | 1 | NA |
| HD003 | MSI | CRC | hereditary | 53 | female | mutated | 1 | NA |
| HD004 | MSI | CRC | hereditary | 42 | male | wildtype | 2 | NA |
| HD005 | MSI | CRC | hereditary | 44 | male | wildtype | 1 | NA |
| HD006 | MSI | CRC | hereditary | 42 | female | mutated | 2 | NA |
| HD007 | MSI | CRC | hereditary | 41 | female | NA | 1 | NA |
| HD008 | MSI | CRC | hereditary | 42 | female | wildtype | 1 | NA |
| HD009 | MSI | CRC | hereditary | 24 | male | NA | 2 | NA |
| HD010 | MSI | CRC | hereditary | 36 | male | wildtype | NA | NA |
| HD011 | MSI | CRC | hereditary | 45 | female | NA | 2 | NA |
| HD012 | MSI | CRC | hereditary | 44 | male | wildtype | NA | NA |
| HD013 | MSI | CRC | hereditary | NA | male | mutated | 3 | NA |
| HD014 | MSI | CRC | sporadic | 72 | male | wildtype | NA | NA |
| HD015 | MSI | CRC | hereditary | NA | male | wildtype | NA | NA |
| HD016 | MSI | CRC | sporadic | 84 | female | wildtype | NA | NA |
| HD017 | MSI | CRC | hereditary | 55 | male | wildtype | NA | NA |
| HD018 | MSI | CRC | hereditary | 45 | male | wildtype | 2 | NA |
| HD019 | MSI | CRC | hereditary | NA | female | NA | NA | NA |
| HD020 | MSI | CRC | hereditary | 41 | female | wildtype | 2 | NA |
| HD021 | MSI | CRC | hereditary | NA | female | wildtype | 2 | NA |
| HD022 | MSI | CRC | sporadic | NA | female | NA | NA | NA |
| HD023 | MSI | CRC | hereditary | NA | female | wildtype | NA | NA |
| HD024 | MSI | CRC | hereditary | 71 | female | wildtype | 1 | NA |
| HD025 | MSI | CRC | hereditary | 62 | male | mutated | 1 | NA |
| HD026 | MSI | CRC | hereditary | 43 | female | wildtype | 1 | NA |
| HD027 | MSI | CRC | hereditary | 71 | male | wildtype | 1 | NA |
| HD028 | MSI | CRC | hereditary | 69 | female | wildtype | 1 | NA |
| HD029 | MSI | CRC | hereditary | 71 | male | wildtype | 1 | NA |
| HD030 | MSI | CRC | sporadic | 53 | female | wildtype | NA | NA |
| HD031 | MSI | CRC | sporadic | 59 | female | wildtype | NA | NA |
| HD032 | MSI | CRC | hereditary | 50 | male | wildtype | NA | NA |
| HD033 | MSI | CRC | hereditary | 48 | male | wildtype | NA | NA |
| HD034 | MSI | CRC | sporadic | 68 | female | wildtype | 1 | NA |
| HD035 | MSI | CRC | sporadic | 74 | male | wildtype | NA | NA |
| HD036 | MSI | CRC | sporadic | 71 | male | wildtype | 3 | NA |
| HD037 | MSI | CRC | sporadic | 75 | female | wildtype | NA | NA |
| HD038 | MSI | CRC | sporadic | 63 | female | wildtype | NA | NA |
| HD039 | MSI | CRC | sporadic | 73 | female | mutated | NA | NA |
| HD040 | MSI | CRC | sporadic | 63 | female | mutated | NA | NA |
| HD041 | MSI | CRC | hereditary | NA | male | NA | 2 | NA |
| HD042 | MSI | CRC | hereditary | NA | male | wildtype | NA | NA |
| HD043 | MSI | CRC | hereditary | NA | male | wildtype | NA | NA |
| HD044 | MSI | CRC | hereditary | 58 | NA | wildtype | 2 | NA |
| HD045 | MSI | CRC | hereditary | NA | female | wildtype | 2 | NA |

| TU ID | MSI.Status | Tumor.type | Etiology (hereditary/sporadic) | age | sex | B2M.Seq | UICC | FIGO |
|---|---|---|---|---|---|---|---|---|
| HD046 | MSI | CRC | hereditary | 43 | female | mutated | 3 | NA |
| HD047 | MSI | CRC | hereditary | 51 | female | mutated | 2 | NA |
| HD048 | MSI | CRC | hereditary | 42 | female | wildtype | 2 | NA |
| HD049 | MSI | CRC | hereditary | 57 | female | wildtype | 2 | NA |
| HD050 | MSI | CRC | hereditary | 36 | female | NA | 3 | NA |
| HD051 | MSI | CRC | hereditary | 54 | female | wildtype | 1 | NA |
| HD052 | MSI | CRC | hereditary | 55 | male | mutated | 1 | NA |
| HD053 | MSI | CRC | hereditary | 33 | male | mutated | 3 | NA |
| HD054 | MSI | CRC | hereditary | NA | female | wildtype | 3 | NA |
| HD055 | MSI | CRC | sporadic | 69 | male | mutated | 2 | NA |
| HD056 | MSI | CRC | sporadic | 55 | female | wildtype | NA | NA |
| HD057 | MSI | CRC | hereditary | 50 | male | mutated | 3 | NA |
| HD058 | MSI | CRC | hereditary | NA | female | NA | NA | NA |
| HD059 | MSI | CRC | hereditary | 72 | male | wildtype | NA | NA |
| HD060 | MSI | CRC | sporadic | 64 | female | wildtype | 1 | NA |
| HD061 | MSI | CRC | hereditary | 49 | male | wildtype | 2 | NA |
| HD062 | MSI | CRC | sporadic | 56 | male | NA | NA | NA |
| HD063 | MSI | CRC | sporadic | 77 | female | wildtype | 2 | NA |
| HD064 | MSI | CRC | hereditary | 44 | male | wildtype | 4 | NA |
| HD065 | MSI | CRC | sporadic | 75 | female | wildtype | 3 | NA |
| HD066 | MSI | CRC | hereditary | 70 | male | wildtype | 2 | NA |
| HD067 | MSI | CRC | sporadic | 70 | male | mutated | 3 | NA |
| HD068 | MSI | CRC | sporadic | 86 | female | wildtype | NA | NA |
| HD069 | MSI | CRC | hereditary | 35 | male | mutated | NA | NA |
| HD070 | MSI | CRC | hereditary | 78 | male | wildtype | 2 | NA |
| HD071 | MSI | CRC | NA | 71 | female | wildtype | NA | NA |
| HD072 | MSI | CRC | sporadic | 72 | female | wildtype | 2 | NA |
| HD073 | MSI | CRC | sporadic | 80 | female | wildtype | NA | NA |
| HD074 | MSI | CRC | sporadic | 76 | female | mutated | 3 | NA |
| HD075 | MSI | CRC | sporadic | 84 | female | wildtype | 3 | NA |
| HD076 | MSI | CRC | sporadic | 81 | female | wildtype | 1 | NA |
| HD077 | MSI | CRC | sporadic | 68 | male | wildtype | NA | NA |
| HD078 | MSI | CRC | hereditary | 39 | male | wildtype | 2 | NA |
| HD079 | MSI | CRC | sporadic | 86 | female | wildtype | 2 | NA |
| HD080 | MSI | CRC | sporadic | 81 | female | wildtype | NA | NA |
| HD081 | MSI | CRC | sporadic | 81 | female | wildtype | 2 | NA |
| HD082 | MSI | CRC | sporadic | 58 | female | mutated | 3 | NA |
| HD083 | MSI | CRC | sporadic | 70 | male | wildtype | 4 | NA |
| HD084 | MSI | CRC | sporadic | 54 | female | mutated | NA | NA |
| HD085 | MSI | CRC | sporadic | 67 | female | mutated | 4 | NA |
| HD086 | MSI | CRC | sporadic | 54 | female | wildtype | NA | NA |
| HD087 | MSI | CRC | hereditary | 55 | male | wildtype | 1 | NA |
| HD088 | MSI | CRC | sporadic | 58 | female | wildtype | 4 | NA |
| HD089 | MSI | CRC | sporadic | 83 | female | wildtype | 2 | NA |
| HD090 | MSI | CRC | sporadic | 84 | male | mutated | 2 | NA |

| TU ID | MSI.Status | Tumor.type | Etiology (hereditary/sporadic) | age | sex | B2M.Seq | UICC | FIGO |
|---|---|---|---|---|---|---|---|---|
| HD091 | MSI | CRC | hereditary | 73 | male | wildtype | 2 | NA |
| HD092 | MSI | CRC | sporadic | 72 | male | wildtype | 2 | NA |
| HD093 | MSI | CRC | sporadic | 65 | female | wildtype | 2 | NA |
| HD094 | MSI | CRC | sporadic | 76 | female | wildtype | 3 | NA |
| HD095 | MSI | CRC | hereditary | 53 | male | wildtype | 2 | NA |
| HD096 | MSI | CRC | sporadic | 76 | female | mutated | 3 | NA |
| HD097 | MSI | CRC | sporadic | 88 | female | wildtype | 2 | NA |
| HD098 | MSI | CRC | hereditary | 55 | male | wildtype | 2 | NA |
| HD099 | MSI | CRC | sporadic | 71 | male | wildtype | 3 | NA |
| HD100 | MSI | CRC | hereditary | 69 | female | wildtype | NA | NA |
| HD101 | MSI | CRC | sporadic | 51 | male | mutated | 3 | NA |
| HD102 | MSI | CRC | hereditary | 40 | male | wildtype | 2 | NA |
| HD103 | MSI | CRC | sporadic | 77 | female | wildtype | 1 | NA |
| HD104 | MSI | CRC | sporadic | 80 | female | wildtype | 3 | NA |
| HD105 | MSI | CRC | sporadic | 75 | male | mutated | 1 | NA |
| HD106 | MSI | CRC | sporadic | 67 | female | wildtype | 1 | NA |
| HD107 | MSI | CRC | hereditary | 39 | male | mutated | 2 | NA |
| HD108 | MSI | CRC | sporadic | 96 | female | wildtype | 2 | NA |
| HD109 | MSI | CRC | hereditary | 27 | male | wildtype | 3 | NA |
| HD110 | MSI | CRC | sporadic | 71 | male | wildtype | 2 | NA |
| HD111 | MSI | CRC | sporadic | 63 | female | wildtype | 2 | NA |
| HD112 | MSI | CRC | sporadic | 71 | male | wildtype | 1 | NA |
| HD113 | MSI | CRC | hereditary | 82 | male | wildtype | 3 | NA |
| HD114 | MSI | CRC | hereditary | 58 | male | wildtype | 2 | NA |
| HD115 | MSI | CRC | sporadic | NA | female | NA | 3 | NA |
| HD116 | MSI | CRC | hereditary | NA | female | wildtype | 2 | NA |
| HD117 | MSI | CRC | hereditary | 67 | male | mutated | 1 | NA |
| HD118 | MSI | CRC | hereditary | 52 | male | wildtype | 1 | NA |
| HD119 | MSI | CRC | hereditary | 59 | male | wildtype | 2 | NA |
| HD120 | MSI | CRC | hereditary | NA | NA | NA | NA | NA |
| HD121 | MSI | CRC | hereditary | NA | NA | NA | NA | NA |
| HD122 | MSI | CRC | hereditary | NA | female | wildtype | NA | NA |
| HD123 | MSI | CRC | hereditary | NA | NA | mutated | NA | NA |
| HD124 | MSI | CRC | hereditary | NA | male | wildtype | NA | NA |
| HD125 | MSI | CRC | hereditary | NA | male | NA | NA | NA |
| HD126 | MSI | CRC | hereditary | 62 | female | NA | NA | NA |
| HD127 | MSI | CRC | hereditary | 59 | female | NA | NA | NA |
| HD128 | MSI | CRC | hereditary | NA | female | wildtype | 2 | NA |
| HD129 | MSI | CRC | hereditary | NA | female | wildtype | 1 | NA |
| HD130 | MSI | CRC | hereditary | NA | male | wildtype | 3 | NA |
| HD131 | MSI | CRC | hereditary | NA | male | wildtype | 1 | NA |
| HD132 | MSI | CRC | hereditary | NA | male | NA | 3 | NA |
| HD133 | MSI | CRC | hereditary | NA | female | wildtype | 4 | NA |
| HD134 | MSI | CRC | hereditary | NA | female | mutated | 2 | NA |
| HD135 | MSI | CRC | hereditary | NA | male | wildtype | 1 | NA |

| TU ID | MSI.Status | Tumor.type | Etiology (hereditary/sporadic) | age | sex | B2M.Seq | UICC | FIGO |
|---|---|---|---|---|---|---|---|---|
| HD136 | MSI | CRC | hereditary | NA | female | wildtype | 2 | NA |
| HD137 | MSI | CRC | hereditary | NA | male | wildtype | 3 | NA |
| HD138 | MSI | CRC | NA | 74 | female | NA | 4 | NA |
| HD139 | MSI | CRC | sporadic | 71 | female | NA | 3 | NA |
| HD140 | MSI | EC | NA | 69 | female | NA | NA | IA |
| HD141 | MSI | EC | NA | 59 | female | NA | NA | IB |
| HD142 | MSI | EC | NA | 81 | female | NA | NA | IA |
| HD143 | MSI | EC | NA | 82 | female | NA | NA | IA |
| HD144 | MSI | EC | NA | 80 | female | NA | NA | II |
| HD145 | MSI | EC | NA | 80 | female | NA | NA | IA |
| HD146 | MSI | EC | NA | 89 | female | NA | NA | IA |
| HD147 | MSI | EC | NA | 84 | female | NA | NA | IIIC |
| HD148 | MSI | EC | NA | 71 | female | NA | NA | IIIA |
| HD149 | MSI | EC | NA | 50 | female | NA | NA | IA |
| HD150 | MSI | EC | NA | 77 | female | NA | NA | IA |
| HD151 | MSI | EC | NA | 68 | female | NA | NA | IB |
| HD152 | MSI | EC | NA | 46 | female | NA | NA | IA |
| HD153 | MSI | EC | NA | 71 | female | NA | NA | IIIC |
| HD154 | MSI | EC | NA | 53 | female | NA | NA | IA |
| HD155 | MSI | EC | NA | 79 | female | NA | NA | IB |

**Table S2. Sample cohort: Overview of sample cohort.** All tumors used in this study were MSI. Tumor type (CRC or EC), etiology (hereditary or sporadic), age, sex, *B2M* sequencing status and stage (UICC for CRC and FIGO for EC) are specified.

| gene name | type | length | %mut | %wt<0.5 | m4 | m3 | m2 | m1 | wt | p1 | p2 | p3 | p4 | %mut | %wt<0.5 | m4 | m3 | m2 | m1 | wt | p1 | p2 | p3 | p4 |
|---|---|---|---|---|---|---|---|---|---|---|---|---|---|---|---|---|---|---|---|---|---|---|---|---|
| BANP | T | 12 | 0,88 | 0,62 | | 0,09 | 0,23 | 0,24 | 0,42 | 0,01 | | | | 1,00 | 0,29 | | | 0,12 | 0,28 | 0,58 | 0,01 | | | |
| ASTE1 | A | 11 | 0,86 | 0,46 | | 0,03 | 0,15 | 0,33 | 0,47 | 0,02 | | | | 0,93 | 0,57 | | | 0,03 | 0,47 | 0,47 | 0,03 | | | |
| CEP164 | A | 11 | 0,78 | 0,21 | | | 0,02 | 0,37 | 0,59 | 0,02 | | | | 0,36 | 0,00 | | | | 0,35 | 0,65 | | | | |
| LTN1 | A | 11 | 0,83 | 0,30 | | 0,01 | 0,13 | 0,30 | 0,55 | | | | | 0,77 | 0,00 | | | | 0,28 | 0,72 | | | | |
| MARCKS | A | 11 | 0,80 | 0,30 | | | 0,05 | 0,36 | 0,56 | 0,02 | | | | 0,54 | 0,08 | | | 0,03 | 0,32 | 0,65 | | | | |
| PTHLH | A | 11 | 0,74 | 0,34 | | 0,01 | 0,12 | 0,34 | 0,52 | 0,01 | | | | 0,82 | 0,36 | | | 0,08 | 0,35 | 0,57 | | | | |
| SLC22A9 | A | 11 | 0,76 | 0,24 | | 0,01 | 0,08 | 0,32 | 0,58 | 0,01 | | | | 0,54 | 0,15 | | | 0,03 | 0,35 | 0,62 | | | | |
| TAF1B | A | 11 | 0,75 | 0,06 | | 0,03 | 0,11 | 0,18 | 0,66 | 0,02 | | | | 0,50 | 0,00 | | | 0,05 | 0,24 | 0,71 | | | | |
| ABCF1 | A | 10 | 0,48 | 0,00 | | | 0,03 | 0,24 | 0,72 | 0,01 | | | | 0,79 | 0,00 | | | | 0,29 | 0,71 | | | | |
| AIM2 | A | 10 | 0,82 | 0,30 | | 0,01 | 0,08 | 0,36 | 0,53 | 0,02 | | | | 0,71 | 0,14 | | | | 0,20 | 0,70 | 0,11 | | | |
| CASP5 | A | 10 | 0,63 | 0,19 | | | 0,02 | 0,35 | 0,59 | 0,04 | | | | 0,92 | 0,83 | | 0,01 | 0,29 | 0,37 | 0,31 | | | | |
| LINC01587 | T | 10 | 0,54 | 0,11 | | | 0,04 | 0,26 | 0,63 | 0,08 | | | | 0,93 | 0,29 | | | | 0,41 | 0,53 | 0,06 | | | |
| RFC3 | A | 10 | 0,43 | 0,07 | | | 0,01 | 0,24 | 0,65 | 0,09 | | | | 0,86 | 0,07 | | | | 0,37 | 0,63 | | | | |
| SLC35F5 | T | 10 | 0,67 | 0,27 | | | 0,06 | 0,40 | 0,53 | 0,01 | | | | 0,71 | 0,07 | | | | 0,32 | 0,68 | | | | |
| SMAP1 | A | 10 | 0,52 | 0,14 | | 0,01 | 0,08 | 0,27 | 0,63 | 0,03 | | | | 0,46 | 0,08 | | | | 0,28 | 0,72 | | | | |
| SPINK5 | A | 10 | 0,26 | 0,01 | | | | 0,28 | 0,70 | 0,02 | | | | 0,08 | 0,00 | | | | 0,16 | 0,84 | | | | |
| TFAM | A | 10 | 0,47 | 0,07 | | 0,02 | 0,07 | 0,25 | 0,65 | 0,01 | | | | 0,38 | 0,00 | | | | 0,31 | 0,69 | | | | |
| TGFBR2 | A | 10 | 0,88 | 0,41 | | 0,01 | 0,08 | 0,41 | 0,47 | 0,03 | | | | 0,54 | 0,15 | | | | 0,38 | 0,62 | | | | |
| TMEM97 | A | 10 | 0,27 | 0,01 | | | | 0,33 | 0,67 | | | | | 0,31 | 0,00 | | | | 0,25 | 0,75 | | | | |
| ELAVL3 | G | 9 | 0,46 | 0,06 | | | 0,01 | 0,20 | 0,66 | 0,13 | | | | 0,31 | 0,00 | | | | 0,23 | 0,77 | | | | |
| EPHB2 | A | 9 | 0,34 | 0,00 | | | | 0,23 | 0,72 | 0,05 | | | | 0,29 | 0,00 | | 0,06 | | 0,18 | 0,76 | | | | |
| FLT3LG | C | 9 | 0,26 | 0,05 | | | | 0,22 | 0,65 | 0,13 | | | | 0,08 | 0,00 | | | | | 0,76 | 0,24 | | | |
| LMAN1 | A | 9 | 0,52 | 0,07 | | | 0,01 | 0,31 | 0,66 | 0,02 | | | | 0,38 | 0,00 | | | 0,06 | 0,27 | 0,67 | | | | |
| NDUFC2 | T | 9 | 0,71 | 0,03 | | | 0,03 | 0,25 | 0,70 | 0,02 | | | | 0,93 | 0,14 | | | 0,02 | 0,38 | 0,57 | 0,02 | | | |
| RGS12 | A | 9 | 0,19 | 0,01 | | | 0,02 | 0,32 | 0,67 | | | | | 0,14 | 0,00 | | | | 0,30 | 0,70 | | | | |
| RUFY2 | A | 9 | 0,16 | 0,00 | | | | 0,24 | 0,70 | 0,06 | | | | 0,21 | 0,00 | | | | 0,36 | 0,64 | | | | |
| TCF7L2 | A | 9 | 0,54 | 0,15 | | | 0,01 | 0,40 | 0,58 | 0,01 | | | | 0,14 | 0,00 | | | | 0,27 | 0,73 | | | | |
| TTK | A | 9 | 0,50 | 0,07 | | | 0,04 | 0,30 | 0,65 | 0,01 | | | | 0,50 | 0,00 | | | | 0,22 | 0,78 | | | | |
| ACVR2A | A | 8 | 0,91 | 0,46 | | 0,01 | 0,08 | 0,36 | 0,53 | 0,02 | | | | 0,29 | 0,00 | | | | 0,25 | 0,75 | | | | |
| BAX | G | 8 | 0,49 | 0,14 | | | 0,00 | 0,20 | 0,73 | 0,06 | | | | 0,29 | 0,07 | | | | 0,34 | 0,66 | | | | |
| GLYR1 | G | 8 | 0,47 | 0,12 | | | | 0,24 | 0,61 | 0,14 | 0,01 | | | 0,14 | 0,07 | | | | 0,37 | 0,52 | 0,11 | | | |
| HNF1A | C | 8 | 0,23 | 0,03 | | | | 0,32 | 0,63 | 0,05 | | | | 0,14 | 0,00 | | | | 0,25 | 0,75 | | | | |
| HPS1 | C | 8 | 0,69 | 0,11 | | | 0,01 | 0,36 | 0,62 | | 0,01 | | | 0,82 | 0,27 | | | | 0,41 | 0,59 | | | | |
| MAPRE3 | C | 8 | 0,32 | 0,00 | | | | 0,30 | 0,68 | 0,02 | | | | 0,00 | 0,00 | | | | | 1,00 | | | | |
| MYH11 | C | 8 | 0,40 | 0,09 | | | 0,02 | 0,34 | 0,58 | 0,06 | | | | 0,29 | 0,00 | | | | 0,37 | 0,63 | | | | |
| OR51E2 | T | 8 | 0,18 | 0,04 | | | | 0,37 | 0,63 | | | | | 0,07 | 0,00 | | | | 0,16 | 0,84 | | | | |
| SRRT | G | 8 | 0,42 | 0,04 | | | 0,01 | 0,24 | 0,69 | 0,05 | 0,01 | | | 0,50 | 0,00 | | | | 0,24 | 0,76 | | | | |
| TFE3 | G | 8 | 0,11 | 0,01 | | | | 0,21 | 0,59 | 0,20 | | | | 0,14 | 0,00 | | | | 0,30 | 0,70 | | | | |
| WASF3 | C | 8 | 0,00 | 0,00 | | | | | 1,00 | | | | | | | | | | | | | | | |
| RNF43.G7 | G | 7 | 0,46 | 0,16 | | | 0,02 | 0,39 | 0,57 | 0,02 | 0,01 | | | | | | | | | | | | | |
| RNF43.C6 | C | 6 | 0,14 | 0,01 | | | | 0,22 | 0,66 | 0,12 | | | | | | | | | | | | | | |

**Table S4. Mutation frequencies and mean allele ratios resulting from ReFrame analysis.** A comprehensive overview of all analyzed cMS, showing the mutation frequencies (%mut), the ratio of samples with biallelic hits, indicated by a wt ratio <0.5 (%wt<0.5), as well as the mean mutational pattern for the cMS candidates sorted by their length. The allele ratios are depicted for wild-type (wt), minus one up to four base pair deletions (m1 – m4) and plus one up to four base pair insertions (p1 – p4).

| Gene | Type | Length | M1>M2 | M2>M1 | M1EXP | M2EXP | Binomial test: P two tailed |
|---|---|---|---|---|---|---|---|
| ABCF1 | A | 10 | 30 | 4 | 27,04 | 6,96 | 0,2872 |
| ACVR2A | A | 8 | 114 | 5 | 94,64 | 24,36 | < 0,0001 |
| AIM2 | A | 10 | 90 | 18 | 85,89 | 22,11 | 0,4033 |
| ASTE1 | A | 11 | 72 | 30 | 81,12 | 20,88 | 0,0359 |
| BANP | T | 12 | 51 | 48 | 78,73 | 20,27 | < 0,0001 |
| BAX | G | 8 | 28 | 8 | 28,63 | 7,37 | 0,836 |
| CASP5 | A | 10 | 63 | 13 | 60,44 | 15,56 | 0,5697 |
| CEP164 | A | 11 | 54 | 5 | 46,92 | 12,08 | 0,0226 |
| ELAVL3 | G | 9 | 32 | 26 | 46,13 | 11,87 | < 0,0001 |
| EPHB2 | A | 9 | 19 | 5 | 19,09 | 4,91 | > 0,9999 |
| FLT3LG | C | 9 | 12 | 7 | 15,11 | 3,89 | 0,0881 |
| GLYR1 | G | 8 | 38 | 19 | 45,33 | 11,67 | 0,0211 |
| HNF1A | C | 8 | 15 | 3 | 14,31 | 3,69 | > 0,9999 |
| HPS1 | C | 8 | 48 | 0 | 38,17 | 9,83 | < 0,0001 |
| LINC01587 | T | 10 | 49 | 16 | 51,69 | 13,31 | 0,4412 |
| LMAN1 | A | 9 | 61 | 6 | 53,28 | 13,72 | 0,0154 |
| LTN1 | A | 11 | 79 | 32 | 88,27 | 22,73 | 0,034 |
| MAPRE3 | C | 8 | 21 | 2 | 18,29 | 4,71 | 0,2022 |
| MARCKS | A | 11 | 81 | 15 | 76,35 | 19,65 | 0,3106 |
| MYH11 | C | 8 | 34 | 9 | 34,20 | 8,80 | > 0,9999 |
| NDUFC2 | T | 9 | 69 | 11 | 63,62 | 16,38 | 0,1651 |
| OR51E2 | T | 8 | 13 | 0 | 10,34 | 2,66 | 0,0843 |
| PTHLH | A | 11 | 66 | 24 | 71,57 | 18,43 | 0,1508 |
| RFC3 | A | 10 | 33 | 16 | 38,97 | 10,03 | 0,0491 |
| RGS12 | A | 9 | 14 | 0 | 11,13 | 2,87 | 0,0891 |
| RNF43.C6 | C | 6 | 6 | 3 | 7,16 | 1,84 | 0,4016 |
| RNF43.G7 | G | 7 | 34 | 3 | 29,42 | 7,58 | 0,0666 |
| RUFY2 | A | 9 | 9 | 3 | 9,54 | 2,46 | 0,7201 |
| SLC22A9 | A | 11 | 73 | 19 | 73,16 | 18,84 | > 0,9999 |
| SLC35F5 | T | 10 | 74 | 7 | 64,42 | 16,58 | 0,0057 |
| SMAP1 | A | 10 | 25 | 10 | 27,83 | 7,17 | 0,292 |
| SPINK5 | A | 10 | 17 | 1 | 14,31 | 3,69 | 0,1485 |
| SRRT | G | 8 | 24 | 6 | 23,86 | 6,14 | > 0,9999 |
| TAF1B | A | 11 | 52 | 40 | 73,16 | 18,84 | < 0,0001 |
| TCF7L2 | A | 9 | 63 | 0 | 50,10 | 12,90 | < 0,0001 |
| TFAM | A | 10 | 32 | 11 | 34,20 | 8,80 | 0,4482 |
| TFE3 | G | 8 | 4 | 4 | 6,36 | 1,64 | 0,0607 |
| TGFBR2 | A | 10 | 80 | 16 | 76,35 | 19,65 | 0,4473 |
| TMEM97 | A | 10 | 20 | 0 | 15,91 | 4,09 | 0,0219 |
| TTK | A | 9 | 49 | 5 | 42,94 | 11,06 | 0,0421 |
| WASF3 | C | 8 | NA | NA | NA | NA | NA |

**Table S5. cMS mutation frames**. Distribution of cMS mutation reading frames. Corresponding cMS length, nucleotide and gene are specified. Column M1>M2 provides the numbers of tumors, for which a higher percentage of M1 compared to M2 alleles has been observed for each of the listed markers. M2>M1 provides the numbers of tumors, for which a higher percentage of M2 compared to M1 alleles has been observed. Given the actual distribution of M1 and M2 frame mutations, expected values (M1EXP and M2EXP) are compared to the observed values of M1 or M2 dominance by a binomial test. Two tailed p-values mark significant deviations towards M1 or M2 dominance.

| | high affinity | low affinity | very low affinity |
|---|---|---|---|
| HLA-A01:01 | 1.4 | 4.9 | 22.5 |
| HLA-A02:01 | 19.7 | 39.5 | 59.7 |
| HLA-A03:01 | 12.0 | 39.0 | 66.5 |
| HLA-A24:02 | 5.1 | 18.6 | 43.7 |
| HLA-A26:01 | 0.2 | 6.0 | 27.4 |
| HLA-B07:02 | 16.9 | 35.5 | 60.4 |
| HLA-B08:01 | 8.1 | 41.1 | 78.8 |
| HLA-B15:01 | 10.9 | 37.6 | 72.7 |
| HLA-B27:05 | 7.4 | 34.4 | 66.5 |
| HLA-B39:01 | 2.8 | 18.3 | 52.0 |
| HLA-B40:01 | 3.7 | 13.9 | 38.1 |
| HLA-B58:01 | 7.9 | 27.9 | 59.0 |

**Table S6. Candidates per HLA**. For each HLA allele and each affinity class (high-affinity, low-affinity, very low-affinity), the percentage of candidates with at least one predicted epitope for that HLA allele is given.

| Gene | cMS repeat | Size (bp) | Forward Primer [5´ > 3´] | Reverse Primer [5´ > 3´] |
|---|---|---|---|---|
| ACVR2A | A8 | 113 | GTTGCCATTTGAGGAGGAAA | CAGCATGTTTCTGCCAATAATC |
| AIM2 | A10 | 76 | TTCTCCATCCAGGTTATTAAGGC | TTAGACCAGTTGGCTTGAATTG |
| ASTE1 | A11 | 117 | ATATGCCCCCGCTGAAATA | TTGGTGTGTGCAGTGGTTCT |
| BANP | T12 | 126 | TTCTGTGGAAGCTCTGCCTT | TCAAGTCGCATCAGATCCAG |
| C4orf6 | T10 | 98 | CCAGAAGCAAATTCACAAGAC | TTTTGCGTGTTCCTTCCTTC |
| CASP5 | A10 | 141 | CAGAGTTATGTCTTAGGTGAAGG | ACCATGAAGAACATCTTTGCCCAG |
| CLOCK | T9 | 73 | TCATTATGTTTAATTTCAGGCTCTTG | CACATATATTATGCTTCCATCTGTCA |
| ELAVL3 | G9 | 134 | GATGCGACCTGTTATCTCCAG | AGGTTGGTCTTGCTGTCGTC |
| GLYR1 | G8 | 113 | GCCTCCAGAAGCTGTGACTT | ATCACCAACATCCCGTCATT |
| LMAN1 | A9 | 114 | CACCCATGTCAGCTTTGCTA | GGAGGAATTTGAGCACTTTCA |
| MARCKS | A11 | 109 | GACTTCTTCGCCCAAGGC | GCCGCTCAGCTTGAAAGA |
| MYH11 | C8 | 77 | CGGGGATTCTCTCTCTGTTTC | CTGAAGGCATGATACCTGGTG |
| NDUFC2 | T9 | 113 | TGAATTTCAGGTTTGCATCG | AACATTTCACGGTCCCTCAC |
| PTHLH | A11 | 107 | TTTCACTTTCAGTACAGCACTTCTG | GAAGTAACAGGGGACTCTTAAATAATG |
| RFC3 | A10 | 60 | TTTCTTTGTCCACAGACTCCATC | GTTACTTGCAATGGTGCTAATTTC |
| SEC63 | A10 | 104 | AGTAAAGGACCCAAGAAAACTGC | TGCTTTTGTTTCTGTTGCTTTG |
| SLC22A9 | A11 | 142 | GCGCCTACAGTGCCTACTCT | GCATGTGGAGCATTTCACAC |
| SLC35F5 | T10 | 102 | TGTGGGGAAACTTACTGCAA | TCAAGTTTCAAACATCATATGCAA |
| TAF1B | A11 | 137 | ACCCAAATAAAAGCCCTCAAC | CTACTTAAAATTCCATTCCATGTCC |
| TCF7L2 | A9 | 75 | GCCTCTATTCACAGATAACTC | GTTCACCTTGTATGTAGCGAA |
| TFAM | A10 | 204 | CTTTGGAAAAAGAAATCATGGAC | AACTATCCCACTTCTGCCTAACTG |
| TGFBR2 | A10 | 149 | GCTGCTTCTCCAAAGTGCAT | CAGATCTCAGGTCCCACACC |
| TTK | A9 | 123 | TTCTTCATCCTCCAAGACTTTT | GATTTCCACAGGGATTCAAGA |
| ZNF294 | A11 | 142 | AAGCCGAAGAGCTCATTGAA | CAGTTGTTAATTCCCAGCCTTC |
| TMEM 97 | A10 | 95 | TGTTGCGGAGCCCCTAC | AACCACCCTGTAGGCATCTC |
| FLT3LG | C9 | 135 | GGGATGACGTGGTGGTG | GTGATCCAGGGCTTCAGC |
| TFE3 | G8 | 135 | CAGAGCAGCTGGACATTGAG | GAAAGTGCAGGTCCAGAAGG |
| SMAP1 | A10 | 94 | TCAAAACTTTGGGCTGTGTTT | TAAGTGGTTTTGCCGGCTT |
| MSH3 | A8 | 147 | AGATGTGAATCCCCTAATCAAGC | ACTCCCACAATGCCAATAAAAAT |
| C1orf34 | G10 | 145 | AGGGACAGGATAGACTGGGG | ATCTCCCAGTCAAAATCCCA |
| SPINK5 | A10 | 141 | TGAGGCGTTTGTTCACTTTG | TGTCATTGCTCCTTTCTCCTG |
| EPHB2 | A9 | 141 | AACATGCAACTCAAACGACG | TTTTATCCCCCGCAAGAAC |
| CEP 164 | A11 | 91 | GTCAACTTCTGGGGCCATTA | ACTCACCAGCGAACTTTTGG |
| ABCF1 | A10 | 114 | GGCAGAAATACAGCAGGGG | CATCATCCACATCCTTCTTCC |
| TCF1 | C8 | 113 | TGGCCATGGACACGTACAG | GTGGACCTTACTGGGGGAGA |
| HPS1 | C8 | 133 | ATGTTATTACCTGTGGCCTGC | CAATACTCACTGCGGCATCT |
| MAPRE3 | C8 | 123 | CTCTTTTCCTCTGGGCAGTTC | GGGCTGATGGAGGATTCTTC |
| OR51E2 | T8 | 139 | TGCAGTGCTCAACAATACAGT | ACAATAGGAGTGCGAGAGGA |
| PRDM2 | A9 | 149 | ATCCTCTCACATCTGCCCTTA | GTGATGAGTGTCCACCTTTCT |
| RGS12 | A9/C8 | 137 | CCGGCTTTCAAAGAGAGAAGA | ACTGGAAACTAACTGTGCATT |
| RUFY2 | A9 | 126 | GGTTCTTCTCTTTAGGACCCC | AGTACTAACCTCAAGAGATCCCT |
| SRRT | G8 | 113 | GTGGCTATGAGATGCCCTATG | CTGGATAGGCAGGACATGGT |

| Gene | cMS repeat | Size (bp) | Forward Primer [5´ > 3´] | Reverse Primer [5´ > 3´] |
|---|---|---|---|---|
| WASF3 | C8 | 114 | CCCTCAACAGACCTCAGCAG | TACAAGGCATGCTGAGTTACC |

**Table S7. Primer sequences for ReFrame analysis.** The primer sequences for each cMS analyzed by ReFrame are depicted. Both, forward and reverse primer as well as the size of the resulting PCR product are shown.

**All data provided as supplementary files (S1-S5) are available on**
https://github.com/atb-data/neoantigen-landscape-msi

**Data S1. (separate file)**

**cMS peptide sequences.** All amino acid sequences extracted from the Seltarbase for the frameshift peptides (FSPs) resulting from one base pair (m1) or two base pair (m2) deletions are depicted. For this purpose, only cMS with a length of 8 or more base pairs, indicated by i.e. A8, representing eight adenine base pairs, are included. For each cMS, eight wildtype amino acids are included at the N-terminus ("including 8 N-terminal wt aa").

**Data S2. (separate file)**

**Predicted HLA binding peptides per candidate and HLA**. For each candidate *neo*antigen (m1 and m2 for each coding microsatellite) and each HLA allele in this study, the presence or absence of binding peptides for the three affinity classes (high-affinity, low-affinity, very low-affinity) is given, together with the candidate *neo*antigen with the highest number of predicted HLA binding peptides.

**Data S3. (separate file)**

**GELS and IRS in various populations.** The calculations of the general epitope likelihood score (GELS) and Immune relevance score (IRS) for different patient cohorts are depicted. The scores for a US European Caucasian, a German population, a Japanese population, as well as two US Hispanic and US African American populations are calculated for high-affinity, low-affinity and very-low affinity epitopes. The calculations were implemented for each cMS peptide sequence (Supplementary table 3) and the respective HLA types including HLA-A01:01, HLA-A02:01, HLA-A03:01, HLA-A24:02, HLA-A26:01, HLA-B07:02, HLA-B08:01, HLA-B15:01, HLA-B27:05, HLA-B39:01, HLA-B40:01 and HLA-B58:01. The different GELS and IRS columns indicate the probability $p_{binding}$, which is assumed as the probability of at least one epitope being truly presented by the HLA molecule. Importantly, values 0.0 indicate no predicted epitope for the appropriate cMS, while "NA" represents the lack of data from the ReFrame analysis, which was only implemented for 82 peptide sequences.

**Data S4. (separate file)**

**ReFrame and epitope prediction scripts.** All scripts are provided, which were used in the present study.

**Data S5. (separate file)**

**Plots depicting predicted HLA binding peptides for all FSP *neo*antigens.**
